# Ionic Regulation of Mechanosurveillance and Metastasis via the MRTFA/KCNMB1 Axis

**DOI:** 10.64898/2026.01.13.699089

**Authors:** Alexa M Gajda, Mohamed Haloul, Vinay Pai, Keyvan Mollaeian, Khushi J Patel, Raymundo Rodriguez-Lopez, Katie M Beverley, Mark A Sanborn, Kihak Lee, Caitlyn C Castillo, Stephanie M Wilk, Beata M Wolska, Faruk Hossen, Eron N Mendenhall, James C Lee, Irena Levitan, Jalees Rehman, Ekrem Emrah Er

**Affiliations:** College of Medicine, Department of Physiology and Biophysics, UIC, Chicago, IL, USA; College of Medicine, Department of Medicine, Division of Pulmonary, Critical Care, Sleep, and Allergy, UIC, Chicago, IL; College of Medicine, Department of Biochemistry and Molecular Genetics, UIC, Chicago, IL, USA; College of Medicine, Department of Medicine, Division of Cardiology, UIC, Chicago, IL, USA; College of Medicine, Department of Biomedical Engineering, UIC, Chicago, IL, USA

**Keywords:** MRTFA, KCNMB1, cancer, cellular stiffness, ion handling, mechanobiology, mechanosurveillance

## Abstract

Cellular stiffness profoundly impacts cancer metastasis at multiple levels, but mechanisms that regulate cancer cells’ stiffness remain poorly understood. Here, we identified potassium efflux and KCNMB1, an auxiliary subunit of the large conductance potassium efflux (BK) channels, as regulators of cellular stiffness downstream of myocardin related transcription factor A (MRTFA). In primary pericytes, KCNMB1 knockdown increased cellular stiffness, which is consistent with the role of potassium efflux in promoting relaxation during excitation-contraction coupling. In a striking contrast, however, KCNMB1 knockdown decreased cellular stiffness in cancer cells. Softer cancer cells were resistant to NK cell mediated cytotoxicity and the low KCNMB1 expression was associated with worse survival in breast cancer patients. Importantly, pharmacological activation of BK channels reduced metastatic burden in mice and improved lysis of cancer cells by cytotoxic T-lymphocytes. These results highlight the unique ionic regulation of stiffness in cancer cells and point to BK channel agonism as a new therapeutic approach in cancer.

## INTRODUCTION

Cellular stiffness is a dynamically regulated property that allows cells to deform with external force and respond to biochemical and mechanochemical stimuli in the microenvironment.^1–3^ Maintenance and regulation of cellular stiffness and mechanoadaptation play key roles in various physiological and pathological contexts. For example, inflammatory stimuli induce dendritic cell (DC) maturation and the accompanying increase in cortical DC stiffness provides the physical stimulus required for the mechanosensitive priming of T cells.^4^ Stiffness of vascular endothelial cells and the sub-endothelial matrix critically regulate vascular barrier function, permeability, and leukocyte adhesion.^5^ In the context of solid tumors, metastatic cells are generally softer than their benign or normal counterparts^1–3^. Cancer cell softening has a non-linear and context-dependent relationship with cellular invasion, but it universally promotes immune evasion from cytotoxic lymphocytes such as CD8^+^ T-lymphocytes and Natural Killer (NK) cells ^3,6–10^. The ability of softer cells to evade cytotoxic lymphocytes is consistent with the ability of these lymphocytes to get better activated when they encounter stiffer targets.^9,11–14^

Multiple biological processes contribute to cellular stiffness. These include intracellular molecular crowding, osmotic balance, lipid composition of the plasma membrane, microtubules and intermediate filaments, and most prominently the filamentous actin (F-actin) cytoskeleton.^15^ F-actin is made up of globular actin monomers (G-actin) and assembles into branches, bundles, and stress fibers to initiate cell motility, maintain cell and organelle architecture, and enforce cell adhesion.^16^ The overall organization of these actin structures is largely influenced by biochemical and mechanochemical cues, as well as the repertoire of actin isoforms and actin-binding proteins that are expressed in the cells. Among F-actin regulators, myocardin-related transcription factor A (MRTFA) plays a central role in transcriptionally regulating actin and actin bundling gene expression in direct response to cytoplasmic G-actin levels.^16–18^ Accordingly, MRTFA expression increases cortical stiffness of cancer cells in an actin-dependent manner, but the precise mechanism of how MRTFA promotes cell stiffness remain unknown.^9^

MRTFA is expressed by breast cancer cells, and it performs both pro-metastatic and anti-metastatic functions^9,19–22^. MRTFA binds to a variety of other transcription factors to activate numerous transcriptional programs.^23^ Some of these cofactors include serum response factor (SRF), the Hippo-pathway effectors yes-associated protein 1 (YAP), the TEA domain transcription factor 1 (TEAD), and transforming growth factor beta (TGFβ) downstream transcription factors: suppressors of mother against decapentaplegic (SMADs). MRTF interactions with these transcription factors have been implicated in cancer cell motility and metastasis in numerous experimental settings.^20,24–26^ Additionally, all these transcription factors also affect actin cytoskeletal remodeling in different developmental, homeostatic, and pathological settings.^23^

Hence, we set out to identify transcriptional partners of MRTFA and its downstream effectors responsible for generating cortical stiffness with the goal of pharmacologically perturbing them to inhibit cancer progression and metastasis. Our structure-function approach identified SRF as a significant contributor to cancer cell stiffening downstream of MRTFA. By using bulk RNA sequencing, we found that MRTFA/SRF axis promotes the expression of genes involved in ion homeostasis. Functionally, MRTFA/SRF promote intracellular calcium concentrations ([Ca^2+^]_i_) and reduce intracellular potassium concentrations ([K^+^]_i_), and the latter involves MRTFA/SRF mediated expression of calcium-activated potassium channel subfamily M regulatory beta subunit 1 (KCNMB1). KCNMB1 is the auxiliary protein responsible for enhancing Ca^2+^ sensitivity of the large-conductance, calcium-activated potassium (BK) channels (encoded by KCNMA1).^27^ We show that KCNMB1 expression significantly contributes to cellular stiffness. Importantly, knockdown of KCNMB1 and the associated JPH2 sensitizes softens cancer cells and makes them resistant to lysis by natural killer cells, which is consistent with the role of target cell stiffness in sensitizing them to cytotoxic lymphocyte driven lysis. Furthermore, upregulation of BK channel activity by a small molecule agonist, BMS-204352 (also known as Maxipost), improves cancer cell stiffness, activation of CD8^+^ T-lymphocytes and inhibits metastatic colonization in mice. Collectively, these data uncover a previously unrecognized role of MRTFA/SRF in regulation of K^+^ ion transport through KCNMB1 and BK channels, which significantly contributes to actin cytoskeleton organization, cellular stiffness and regulation of metastatic potential in cancer cells.

## RESULTS

### SRF is necessary for MRTFA-mediated cortical cell stiffness

To identify regulators of cellular stiffness, we aimed to narrow down the list of MRTFA transcription partners involved in regulating cellular architecture. For this, we targeted YAP, TGFß, and SRF transcriptional programs in MRTFA-overexpressing E0771 mouse mammary tumor cells (**Figure 1A**). Verteporfin and SB-505124 reduced YAP levels and SMAD2/3 phosphorylation, respectively, but SRF knockdown was the most effective in promoting cell rounding (**Figures 1B, S1A and S1B**). To further probe the role of SRF in MRTFA-driven cellular architecture, we mutated the tyrosine 238 residue within the B1 region of MRTFA to alanine. Y238A mutation in MRTFA abolishes SRF binding and the identical Y305A mutation in MRTFB spares SRF independent transcription factors (**Figure S1C**).^23,26,28^ MRTFA^Y238A^ failed to induce the expression of known SRF target genes despite localizing to the nucleus as well as the endogenous and wildtype MRTFA in mouse mammary tumor cells, AT-3 and E0771 (**Figures 1C and S1D – S1G**). Consistent with the role of MRTFA/SRF in regulating F-actin cytoskeleton, we found that MRTFA^WT^, but not MRTFA^Y238A^ promoted F-actin stress fiber alignment as judged by the increased level of anisotropy (**Figures 1D and 1E**). F-actin stress fiber anisotropy correlates with cortical stiffness as measured by atomic force microscopy (AFM) in cultured cells.^29^ Accordingly, MRTFA^WT^ expressing cells were significantly stiffer than MRTFA^Y238A^ expressing cells and they were stiffer than cells that co-expressed an SRF short-hairpin RNA (**Figures 1F, 1G and S1H**). Together, these data show that MRTFA/SRF binding is necessary for inducing F-actin organization and for promoting cancer cell stiffness and validate the use of MRTFA^WT^ and MRTFA^Y238A^ expression as tools for identifying regulators of cellular stiffness.

**Figure 1.**
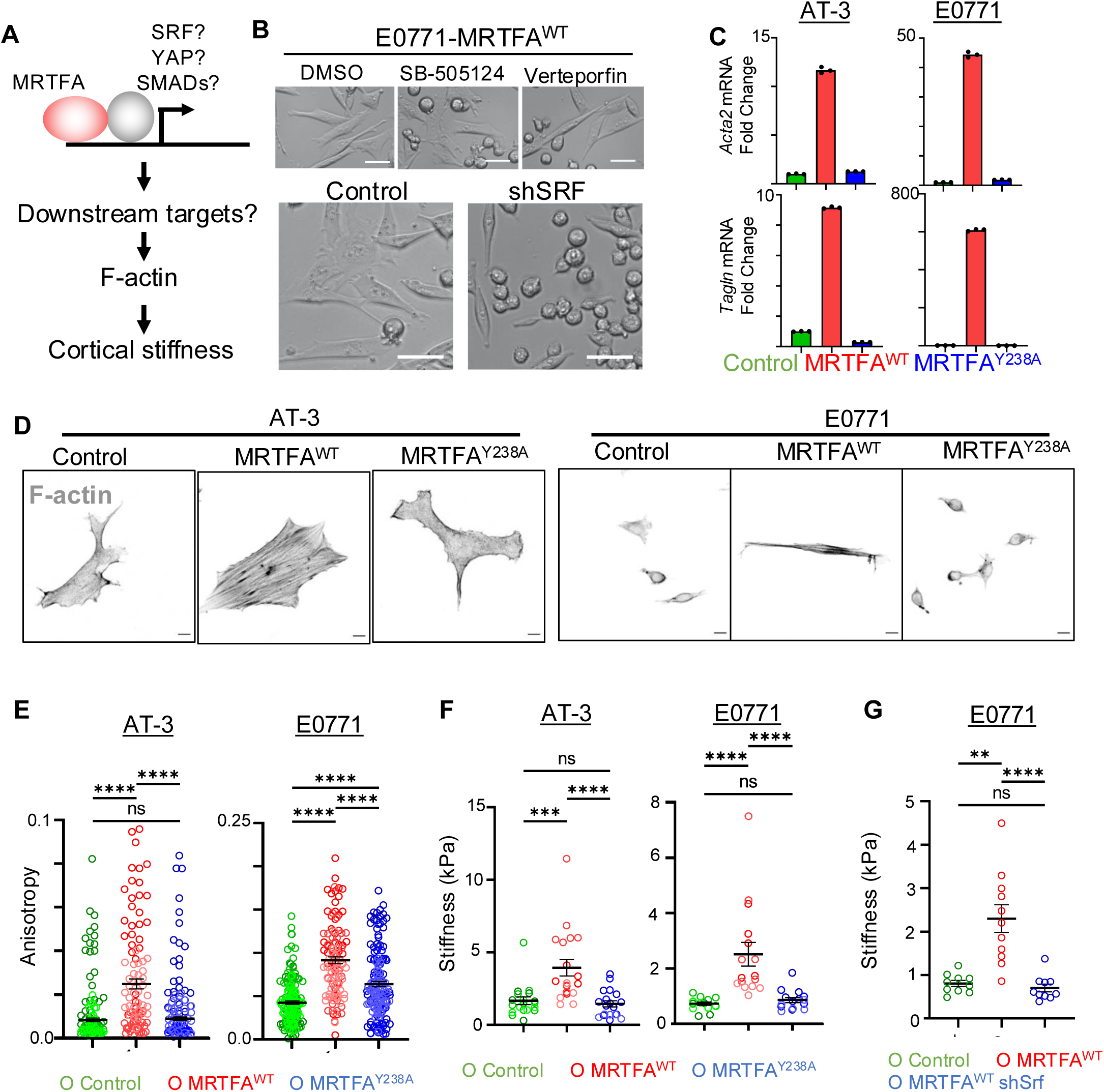
SRF is necessary for MRTFA-mediated cortical cell stiffness. (A) Diagram depicting different MRTFA partners that can promote cortical stiffness in cancer cells. (B) Brightfield images of E0771 cells overexpressing MRTFA^WT^ treated with DMSO (vehicle only control), TGFβ inhibitor (SB-505124; 2.5 µM) or YAP inhibitor (Verteporfin; 10 µM) for 24 hrs. Brightfield images of E0771 cells overexpressing MRTFA^WT^ in the absence or presence of shRNA mediated SRF knockdown (shSRF). Scale bars: 50 µm. (C) Relative transcript levels of downstream MRTFA target genes, *Acta2* and *Myh11*. Data are shown as fold change calculated from 3 technical replicates per sample and normalized GAPDH. Each graph is representative of 3 independent experiments. (D) Confocal images of representative cells stained with phalloidin (F-actin). Scale bar = 10 µm. (E) Anisotropy measurements for cell lines in (D). Each graph is a compilation of 3 independent experiments; AT-3: pRetroX = 95 cells, MRTFA^WT^ = 95 cells, MRTFA^Y238A^ = 92 cells; E0771: Control (pRetroX) = 165 cells, MRTFA^WT^=112 cells, MRTFA^Y238A^=158 cells; Kruskal-Wallace with multiple comparisons. All error bars are mean±s.e.m. ns, not significant; \**P*<0.05; \*\**P*<0.01; \*\*\**P*<0.001. (F) Raw AFM stiffness measurements comparing MRTFA^WT^ and MRTFA^Y238A^ compared to control cells. Each graph is a compilation of 2 independent experiments. AT-3: pRetroX = 19 cells, MRTFA^WT^ = 19 cells, MRTFA^Y238A^ = 19 cells; E0771: Control (pRetroX) = 15 cells, MRTFA^WT^ = 16 cells, MRTFA^Y238A^ = 15 cells; Kruskal-Wallace with multiple comparisons. All error bars are mean±s.e.m. ns, not significant; \**P*<0.05; \*\**P*<0.01; \*\*\**P*<0.001. (G) Raw AFM stiffness measurements comparing SRF knockdown effect on MRTFA^WT^ cell stiffness. Results are combined from 2 independent experiments. E0771: Control (pRetroX) = 10 cells, MRTFA^WT^ + Control (pLKO) = 11 cells, MRTFA^WT^ + shSRF=10 cells; Kruskal-Wallace with multiple comparisons. All error bars are mean±s.e.m. ns, not significant; \**P*<0.05; \*\**P*<0.01; \*\*\**P*<0.001.

### mRNA sequencing reveals MRTFA/SRF axis as a regulator of monoatomic ion transport genes

We performed bulk mRNA sequencing on MRTFA^WT^ and MRTFA^Y238A^ expressing E0771 and AT-3 cell lines to discover new genes involved in MRTFA/SRF-driven cell stiffness. A total of 949 and 218 genes were significantly upregulated by at least two-fold by MRTFA^WT^ and MRTFA^Y238A^, respectively. We reasoned that the “stiffness genes” would be within the set of genes that were exclusively upregulated by MRTFA^WT^, but not by MRTFA^Y238A^, expression in both cell types (**Figures 2A**). Gene Ontology (GO) terms that were enriched by MRTFA^WT^ expression were mostly related to the actin cytoskeleton, as expected (**Table S1**). GO terms that were enriched by MRTFA^Y238A^ were related to antiviral defense, consistent with the SRF-independent role of MRTFA in regulating histone modifiers, signal transducer and activator of transcription (STAT) signaling, and epigenetic modifications (**Figure S2A and Table S2**).^25,30^

**Figure 2.**
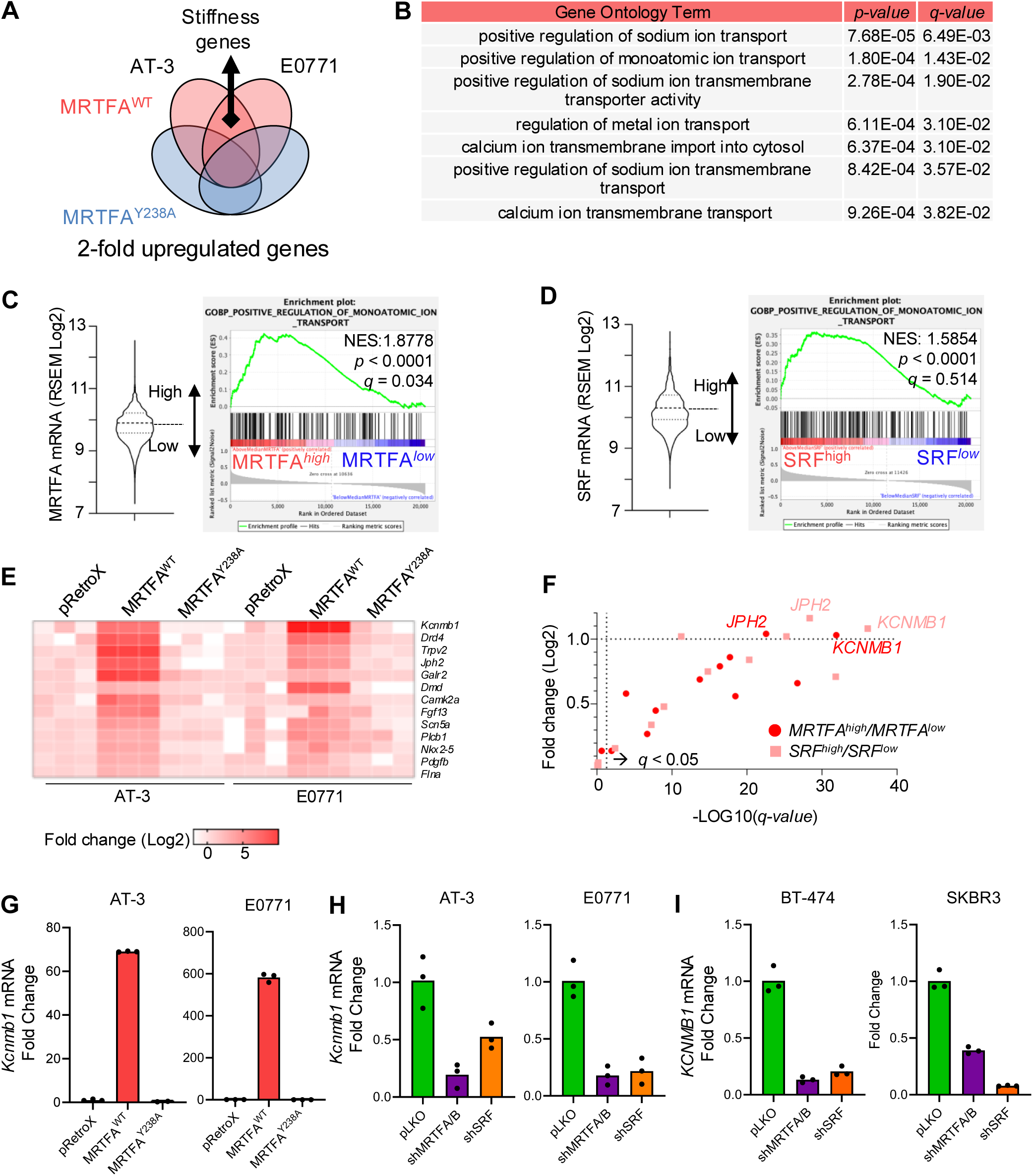
MRTFA/SRF promotes expression of monoatomic ion channel genes. (A) Venn diagram representing the location of the putative stiffness genes in AT-3 and E0771 cell lines where they are exclusively upregulated at least 2-fold by MRTFA^WT^ expression. (B) Gene Ontology analysis of putative “stiffness genes” from A. See Table S1 for full GO term enrichment sets. (C) GSEA analysis performed on patient data from TCGA PanCancer Atlas of invasive breast cancer by using “monoatomic”, “channel” or “transport” keyword containing gene signatures. Patients’ samples were allocated to MRTFA^high^ or MRTFA^low^ categories based on the median MRTFA expression value. (D) Same GSEA analysis as (C), except patients’ samples were allocated to SRF^high^ versus SRF^low^ categories based on the median SRF expression value. (E) Heat map of bulk RNA-seq data showing top ion-associated genes upregulated in AT-3 or E0771 MRTFA^WT^ cells compared to MRTFA^Y238A^. (F) mRNA expression values of all genes in (E) accessed through cBioportal in MRTFA^high/low^ and SRF^high/low^ patients in TCGA. Multiple hypothesis testing correction for statistical significance and the resulting *q* values were calculated by cBioportal. See Table S3 for all genes. (G) Relative transcript levels of *Kcnmb1* in MRTFA^WT^ and MRTFA^Y238A^ cell lines. Data are shown as fold change calculated from 3 technical replicates per sample and normalized to GAPDH. Each graph is representative of 3 independent experiments. (H-I) Relative transcript levels of KCNMB1 in mouse and human shMRTFA/B and shSRF cell lines. Data are shown as fold change calculated from 3 technical replicates per sample and normalized GAPDH. Each graph is representative of 3 independent experiments.

Interestingly, we identified several GO terms involved in monoatomic ion transporter terms in our GO analyses (**Figure 2B**). To determine whether the associations between monoatomic ion transport expression patterns and MRTFA and SRF expression levels were prevalent in human cancer, we interrogated mRNA sequencing data of samples obtained from breast cancer patients in The Cancer Genome Atlas (TCGA). We binned tumor samples into MRTFA^high^, MRTFA^low^, SRF^high^, and SRF^low^ categories based on the median expression of respective genes and used Gene Signature Enrichment Analysis (GSEA) to interrogate gene signatures that contained the terms ‘monoatomic’, ‘channel’ or ‘transport’. Indeed, MRTFA^high^ patient samples showed a statistically significant enrichment in multiple gene signatures, such as “positive regulation of monoatomic ion transport” and “calcium ion transmembrane import into cytosol” (**Figures 2C**, **2D, S2B and S2C**). These terms were also enriched in SRF^high^ samples, but they did not pass a multiple hypothesis testing, likely due to the ability of SRF in partnering with Ternary Complex Factors of the ETS family of transcription factors.^28^

### MRTFA-SRF drives KCNMB1 expression in cancer cells

Since there was an enrichment of ion channel terms in both the MRTFA^WT^ expressing cell lines and the breast cancer patient samples, we investigated whether any of the individual genes upregulated by MRTFA^WT^ were also highly expressed in MRTFA^high^ and SRF^high^ patient samples. Among this set of MRTFA target genes, expression of KCNMB1 was one of the most significantly upregulated genes both in mouse cell lines and in patient samples (**Figures 2E-2F and Table S3**). We further validated MRTFA driven KCNMB1 expression by qPCR (**Figures 2G**). Concordantly, knockdown of endogenous MRTF and SRF in mouse and human breast cancer cell lines, SKBR3 and BT-474, downregulated KCNMB1 expression levels, but perturbing YAP or TGFβ signaling did not robustly influence KCNMB1 expression (**Figures 2H**, **2I, S2D-S2F**). Importantly, our analyses of chromatin immunoprecipitation sequencing (ChIP-Seq) data revealed several experimentally validated SRF binding sites in the promoter and enhancer regions of the mouse and human *KCNMB1* gene loci and we were able to validate the direct interaction between the MRTFA and SRF proteins with the Kcnmb1 enhancer by using ChIP-qPCR (**Figure S2G and S2H**). Exogenous expression of myocardin (MYOCD), a transcription factor related to MRTFA and MRTFB was previously shown to activate the *Kcnmb1* promoter in reporter assays *in vitro*, but we found that MYOCD expression is barely detectable in our cells (**Figure S2I**).^31^ Together, these findings revealed KCNMB1 as a direct transcriptional target of endogenous MRTF-SRF signaling and prompted us to determine how MRTFA/SRF regulate intracellular potassium concentrations.

### MRTFA/SRF is a regulator of intracellular potassium levels

Intracellular potassium concentrations ([K^+^]_i_) reach 150 mM and we reasoned that this high concentration could make it difficult to measure small differences in magnitude.^32^ Therefore, we used a cell-permeable K^+^ dye, IPG-1AM, which has the highest K_D_ among all related dyes.^33^ Overexpression of MRTFA^WT^ lowered [K^+^]_i_ compared to control and to cells that overexpress MRTFA^Y238A^ (**Figure 3A**). Concordantly, MRTFA/B and SRF knockdown significantly elevated [K^+^]_i_ (**Figure 3B**). We also measured intracellular calcium concentrations in cells with MRTFA/SRF perturbations by using a previously described Fluo-4/FuraRed ratiometric calcium imaging method.^34–36^ In these analyses, [Ca^2+^]_i_ increased by MRTFA^WT^ expression, and they were decreased by knockdown of MRTFA and SRF (**Figures S3A-S3B**). Taken together with the results of RNA sequencing experiments, these data reveal the role of MRTFA/SRF in regulation of [K^+^]_i_ and [Ca^2+^]_i_.

**Figure 3.**
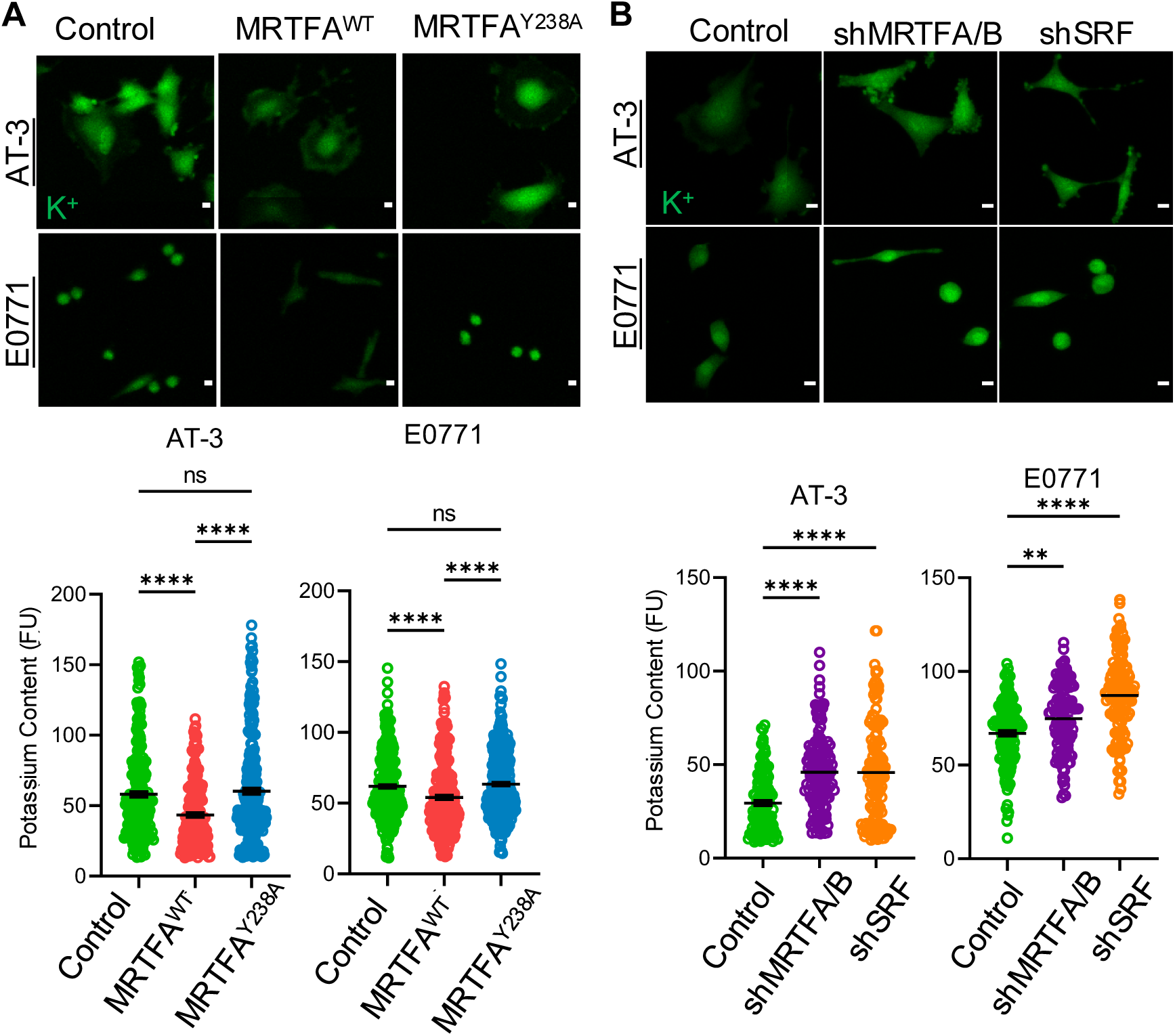
MRTFA/SRF is a regulator of intracellular potassium levels. (A and B) Top: Confocal images of representative cells loaded with IPG-1 AM K^+^ binding dye. Scale bar = 10 µm. Bottom: Graphs showing mean fluorescence intensity of intracellular K^+^ bound dye. Each graph is representative of 3 independent experiments; AT-3: Control (pRetroX) = 261 cells, MRTFA^WT^ = 196 cells, MRTFA^Y238A^ = 291 cells, Control (pLKO) = 125 cells, shMRTFA/B=134 cells, shSRF = 140 cells; E0771: Control (pRetroX) = 353 cells, MRTFA^WT^ = 258 cells, MRTFA^Y238A^ = 362 cells, Control (pLKO) = 147 cells, shMRTFA/B = 119 cells, shSRF = 132 cells; Kruskal-Wallace with multiple comparisons. All error bars are mean±s.e.m. ns, not significant; *P<0.05; **P<0.01; ***P<0.001.

### Potassium homeostasis regulates cancer cell stiffness

Ca^2+^ influx promotes cytoskeletal contractility, and several proteins involved in contractility also promote cellular stiffness.^15^ Accordingly, we found that MRTFA-induced F-actin anisotropy and cellular stiffening were reversed by treatment of cells with BAPTA-AM (1,2-bis(o-aminophenoxy) ethane-N,N,N′,N′-tetraacetic acid) and EDTA (Ethylenediaminetetraacetic acid), which chelate Ca^2+^ within and outside the cells, respectively, without causing cell death or MRTFA mislocalization (**Figures S3C-S3I**). These results confirm the role of cellular Ca^2+^ on contractility and stiffness, but the role of K^+^ in cell stiffness is not well understood.

To determine whether K^+^ regulated cancer cell stiffness, we devised a series of genetic gain-of-function, loss-of-function and epistasis experiments by using MRTFA target gene KCNMB1^WT^, its naturally occurring gain-of-function variant, KCNMB1^E65K^, and Iberiotoxin, a specific non-cell permeable toxin that inhibits BK channels (**Figure 4A**).^37–39^. During electrical excitation-contraction (EC) coupling, an influx of Ca^2+^ promotes calcium-induced calcium release (CICR) from the endo/sarcoplasmic reticulum (ER/SR). This localized Ca^2+^ binds KCNMB1 and enforces its open state and thereby promotes K^+^ efflux. The KCNMB1^E65K^ variant is annotated as the rs11739136 alternate allele in dbSNP and ClinVar databases and it has a lower Ca^2+^ threshold for promoting the open state of BK channels than KCNMB1^WT^. We deployed two orthogonal methods to validate that KCNMB1^E65K^ promotes K^+^ efflux in cancer cells: fluorogenic [K^+^]_i_ measurements and whole-cell patch-clamp to measure outward K^+^ currents as a result of K^+^ efflux. KCNMB1^E65K^ expression significantly reduced [K^+^]_i_ compared to KCNMB1^WT^ and control cells (**Figure 4B**). KCNMB1^E65K^ also produced larger K^+^ currents than KCNMB1^WT^ (**Figure S4A-C**). Importantly, KCNMB1^E65K^ increased F-actin anisotropy and cellular stiffness in comparison to control cells and to KCNMB1^WT^ overexpressing cells, but it did not increase [Ca^2+^]_i_ (**Figures 4C, 4D, S4D**). Treating KCNMB1^E65K^ expressing cancer cells with IbTX was sufficient to reduce F-acting anisotropy and soften cells (**Figures S4E - S4H**). These data suggest that KCNMB1 gain-of-function promotes cell stiffness by upregulation F-actin cytoskeleton polymerization through increased K^+^ efflux.

**Figure 4.**
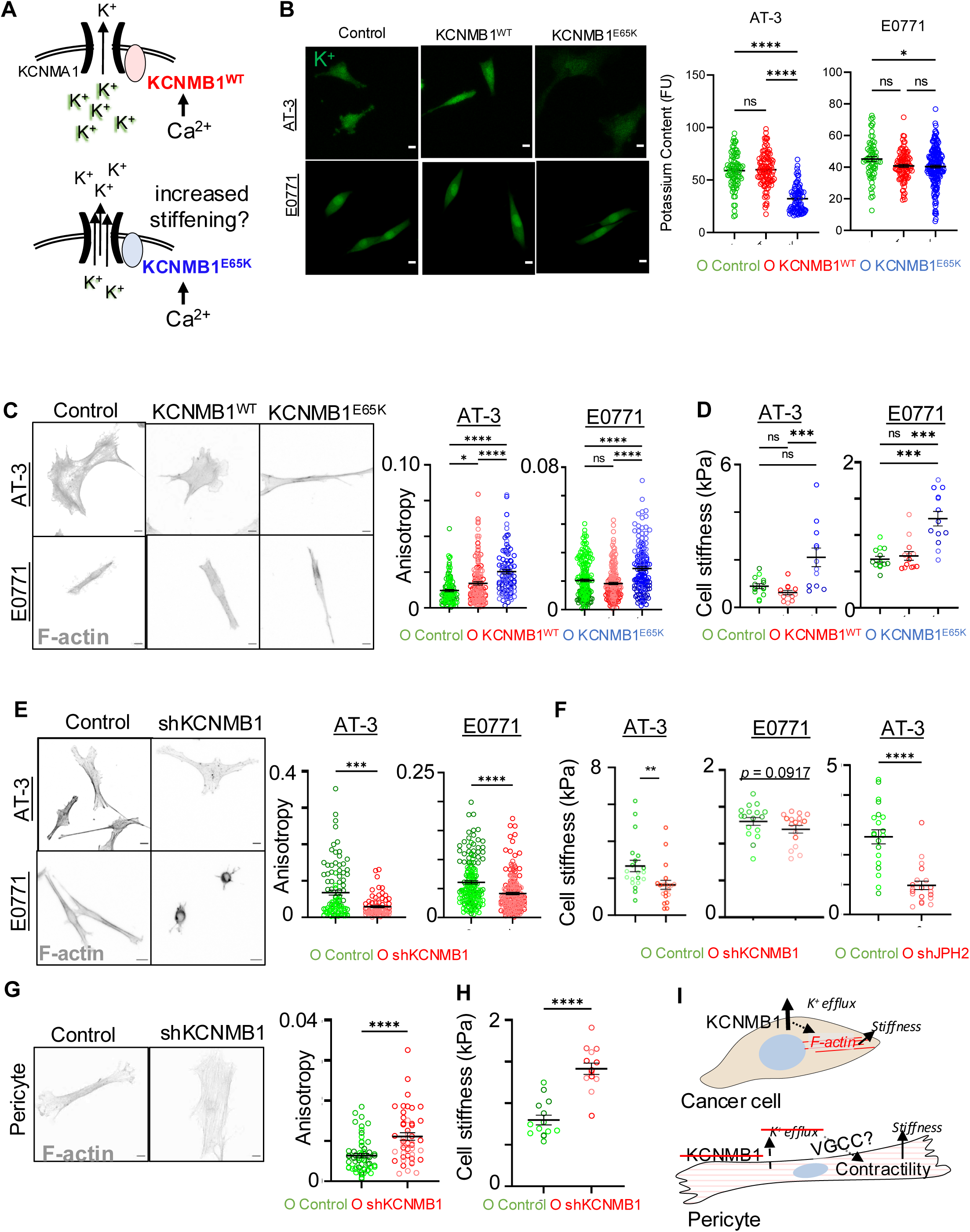
Hyperactive KCNMB1^E65K^ activity promotes cancer cell stiffness. (A) Schematic demonstrating the functional differences in KCNMB1^WT^ and its gain-of-function germline variant KCNMB1^E65K^. For the same amount of intracellular Ca^2+^, KCNMB1^E65K^ promotes a higher degree of K^+^ efflux through the BK channels than the KCNMB1^WT^. KCNMA1 is the pore forming subunit of BK channels. (B) Left: Confocal images of representative cells loaded with IPG-1 AM K^+^ binding dye. Scale bar = 10 µm. Right: Graphs showing mean fluorescence intensity of intracellular K^+^ bound dye. Each graph is representative of 3 independent experiments; AT-3: Control (pRetroX) = 82 cells, KCNMB1^WT^ = 101 cells, KCNMB1^E65K^ = 79 cells; E0771: Control (pRetroX) =63 cells, KCNMB1^WT^ = 96 cells, KCNMB1^E65K^ = 185 cells; Kruskal-Wallace with multiple comparisons. All error bars are mean±s.e.m. ns, not significant; \**P*<0.05; \*\**P*<0.01; \*\*\**P*<0.001. (C) Left: Confocal images of representative AT-3 and E0771 KCNMB1^WT^ and KCNMB1^E65K^ cells compared to controls. Cells are stained with phalloidin (F-actin). Scale bar = 10 µm. Right: Anisotropy measurements for corresponding cell lines. Each graph is a compilation of 3 independent experiments that are represented by different shades of colors. AT-3: Control (pRetroX) = 108 cells, KCNMB1^WT^ = 142 cells, KCNMB1^E65K^ = 95 cells; E0771: Control (pRetroX) = 181 cells, KCNMB1^WT^ = 176 cells, KCNMB1^E65K^ = 156 cells; Mann-Whitney test. All error bars are mean±s.e.m. ns, not significant; \**P*<0.05; \*\**P*<0.01; \*\*\**P*<0.001. (D) AFM stiffness measurements of AT-3 and E0771 KCNMB1^WT^ and KCNMB1^E65K^ cells compared to pRetrox controls. Results are from 3 combined experiments. AT-3: Control (pRetroX) = 14 cells, KCNMB1^WT^ = 13 cells, KCNMB1^E65K^ = 12 cells; E0771: Control (pRetroX) = 13 cells, KCNMB1^WT^ = 13 cells, KCNMB1^E65K^ = 13 cells; Mann-Whitney test. All error bars are mean±s.e.m. ns, not significant; \**P*<0.05; \*\**P*<0.01; \*\*\**P*<0.001. (E) Left: Confocal images of representative AT-3 and E0771 KCNMB1 knockdown cells compared to control. Cells are stained with phalloidin (F-actin). Scale bar=10 µm. Right: Anisotropy measurements for corresponding cell lines. Graphs are representative of 3 independent experiments. AT-3: Control (pLKO) = 45 cells, shKCNMB1 = 29 cells; E0771: Control (pLKO) = 68 cells, shKCNMB1 = 100 cells; Mann-Whitney test. All error bars are mean±s.e.m. ns, not significant; \**P*<0.05; \*\**P*<0.01; \*\*\**P*<0.001. (F) AFM stiffness measurements of AT-3 and E0771 KCNMB1 knockdown and AT-3 JPH2 knockdown cells compared to controls. AT-3 and E0771 graphs are compiled from 2 independent experiments. AT-3: Control (pLKO) = 20 cells, shKCNMB1 = 20 cells; E0771: Control (pLKO) = 17 cells, shKCNMB1 = 17 cells; AT-3: Control (pLKO) = 20 cells, shJPH2 = 20 cells; Mann-Whitney test. All error bars are mean±s.e.m. ns, not significant; *P<0.05; **P<0.01; ***P<0.001. (G) Left: Confocal images of representative pericytes with KCNMB1 knockdown compared to control. Cells are stained with phalloidin (F-actin). Scale bar = 10 µm. Right: Anisotropy measurements for corresponding cell lines. Results are a compilation of 3 independent experiments. Pericytes: Control (pLKO) = 59 cells, shKCNMB1 = 44 cells; Mann-Whitney test. All error bars are mean±s.e.m. ns, not significant; \**P*<0.05; \*\**P*<0.01; \*\*\**P*<0.001. (H) AFM stiffness measurements of representative pericytes with KCNMB1 knockdown compared to control. Results are from 3 combined experiments. Pericytes: pLKO = 14 cells, shKCNMB1 = 14 cells; Mann-Whitney test. All error bars are mean±s.e.m. ns, not significant; \**P*<0.05; \*\**P*<0.01; \*\*\**P*<0.001. (I) Model demonstrating how the role of KCNMB1 in cancer cells contrasts its role in primary pericytes that perform excitation-contraction coupling. KNCMB1 functions to stiffen cancer cells, while it functions to soften pericytes by shutting off voltage gated calcium channels (VGCC).

KCNMB1^WT^ expression alone was not sufficient to promote cell stiffness, which pointed to the existence of other MRTFA/SRF proteins that are required for KCNMB1^WT^ function. Indeed, we identified JPH2 as another MRTFA/SRF-dependent protein both in mouse cancer cell lines and in the human TCGA data (**Figures 2E and 2F**). During EC cycle, JPH2 adjoins ER/SR to KCNMB1 and to the BK channels.^40,41^ Knocking down either JPH2 or KCNMB1 in AT-3 and E0771 cell lines reduced cancer cell stiffness and reduced F-actin anisotropy (**Figure 4E**, **4F, S4I**). These data show that JPH2 and KCNMB1 are significant contributors to stiffening of cancer cells, but they contradict the conventional view that loss of JPH2-BK channel coupling should stiffen cells by increasing cellular contractility. This is because loss of JPH2 promotes Ca^2+^ leak from ER/SR and disrupts K^+^ efflux that would otherwise inhibit voltage-gated calcium influx channels (VGCC) in electrically excitable cells, such as cardiomyocytes and intact arteries. Thus, we measured the biophysical effects of KCNMB1 knockdown on another electrically excitable cell type: primary human brain pericytes. In striking contrast to cancer cells, KCNMB1 knockdown increased F-actin anisotropy, which was coupled with an increase in cellular stiffness (**Figure 4G**, **4H and S4J**). Collectively, these results suggest that KCNMB1 promotes cellular stiffness uniquely in cancer cells, whereas it promotes cellular relaxation and softening in cells that physiologically perform EC coupling and that robustly express VGCCs, such as pericytes (**Figure 4I**).

### Expression of BK channel proteins is associated with better patient outcomes and sensitivity to immune surveillance

Given the unique biology of KCNMB1 and cellular stiffness in cancer cells, we set out to determine the broader functional consequence of BK channel expression in cancer biology. For this, we first measured the correlation between the expression of KCNMB1, JPH2 and the BK channel protein, KCNMA1, and patient survival. Interestingly, expression of all three genes was associated with better patient survival in multiple breast cancer subtypes (**Figures 5A**, **5B and Table S4**). Expression of KCNMA1 was significantly reduced in Basal subtypes of tumors, which are more aggressive, and all three proteins were downregulated in metastatic tumors (**Figures 5C, S5A, S5B and Table S5**). To determine whether this correlation reflected an anti-metastatic role of BK channel proteins, we interrogated the effect of KCNMB1 perturbations on multiple metastatic phenotypes such as proliferation, migration and immune evasion because all these phenotypes can be affected by changes in cellular stiffness in different contexts (**Figure 5D** reviewed in Refs ^1,2,6,7,9,10^). KCNMB1 knockdown or overexpression did not change wound healing or proliferation (**Figures 5E**, **5F, S5C, S5D**). Interestingly, however, time-lapse imaging of co-cultured cancer cells and primary mouse NK cells showed that NK cells took longer to kill softer cancer cells with KCNMB1 and JPH2 knockdowns (**Figures 5G and S5E**).

**Figure 5.**
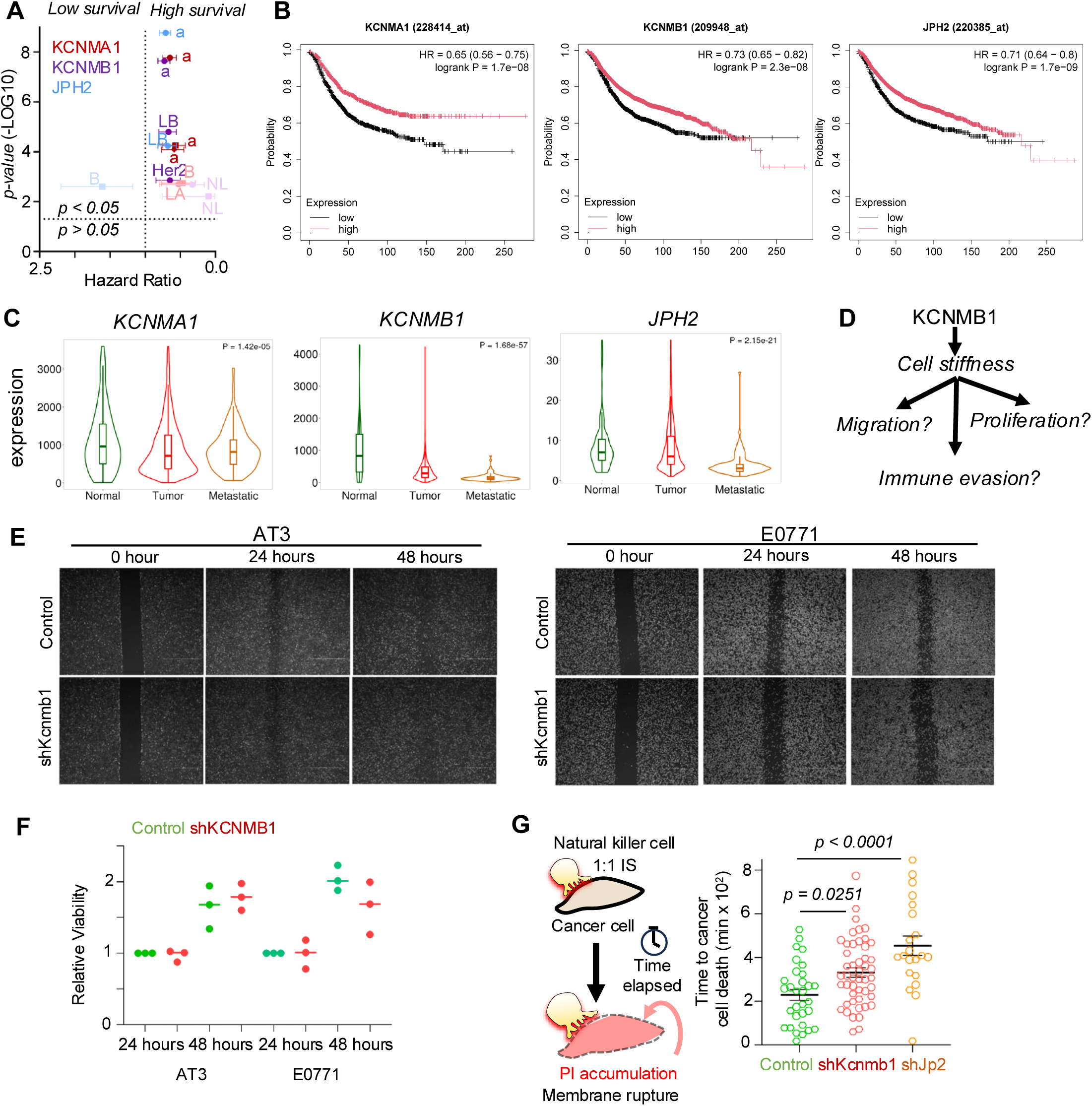
Correlation between expression of BK channel proteins and malignant phenotypes. (A) Association between high KCNMA1 (red), high KCNMB1 (purple) and high JPH2 (blue) mRNA expression and survival in breast cancer patients from all PAM50 subtypes (“a”) or Luminal A (“LA”), Luminal B (“LB”), Normal Like (“NL”), Basal (“B”), or Her2+ (“Her2”). Solid circles (“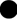”), squares (“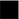”), diamonds (“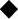”) represent overall survival, distant metastasis free survival, and relapse free survival, respectively. Dark and light colors represent FDR ≤ 1% and FDR = 20%. (B) Examples of Kaplan-Meier plots for overall survival of breast cancer patients expressing high and low KCNMA1, KCNMB1, and JPH2 mRNA in the Kmplotter microarray datasets. Expression thresholds were set automatically, and false discovery rate is less than 1% for all inquiries. (C) Expression of listed genes in normal (non-cancerous), primary tumor and metastasis samples obtained from patients by using TNMplot tool. (D) Schematic of what KCNMB1 mediated cell stiffening could hypothetically regulate. (E) Representative images of wound healing assays to determine whether KCNMB1 knockdown differentially regulates migration and proliferation. Scale bars: 800 µM. Results are representative of 3 independent experiments. (F) Relative viability of control and KCNMB1 knockdown cells by using cell titer glow. Graph shows all three independent experiments performed in triplicate. (G) Schematic representing natural killer and cancer cell co-culture experiments and quantitation of time to cell death following 1:1 contact and immune synapse formation with a natural killer cell. Co-cultures were imaged for 18 hours, and cancer cell death was determined by propidium iodide (PI) accumulation and cellular rupture. Graphs represent data compiled from 2 independent experiments. Control (pLKO) = 30 cells, shKCNMB1 = 46 cells, shJPH2 = 21 cells; Mann-Whitney test. All error bars are mean±s.e.m.

### Pharmacological activation of BK channels hinders metastatic colonization in a cytotoxic lymphocyte dependent manner

Based on clinical data showing a negative association between BK channel gene expression and patient outcomes, we set out to test a clinically feasible pharmacological strategy to reinvigorate the exhausted cytotoxic lymphocytes and to activate BK channels. CD8^+^ T lymphocytes are reinvigorated by anti-PD1 antibody treatment in patients with metastatic breast cancer.^42^ BK channel activation is accomplished by BMS-204352, a small molecule BK channel agonist that has been previously used in human clinical trials to treat stroke patients.^43^ Thus, we treated immune competent mice with a combination of anti-PD1 antibody and BMS-204352 treatment either 10 days after orthotopic implantation into the mammary fat pad or 4 days after metastatic seeding in the lungs, which was accomplished by tail-vein injection. Anti-PD1 antibody treatment was effective in restraining orthotopic tumor growth, and this was not altered by BMS-204352 treatment (**Figures 6A and 6B**). In the metastatic setting however, BMS-204352 treatment alone or in combination with anti-PD1 treatment after the establishment of metastatic colonies strikingly reduced further metastatic colonization by disseminated cancer cells (**Figure 6C**). Granted that both anti-PD1 treatments and K^+^ efflux perturbations can have significant effects on cardiovascular function in patients, we used echocardiography to measure cardiac function in our preclinical model.^44,45^ Results showed no significant differences between mice treated with control and BMS-204352 and anti-PD1 combination in any of the cardiovascular parameters we measured such as stroke volume, ejection fraction and cardiac output (**Figure S6A-S6J**). These data suggest that reinvigorating T-lymphocytes by anti-PD1 treatment improves the anti-metastatic effect of BMS-204352, and this drug combination is safe from a cardiovascular perspective.

**Fig. 6.**
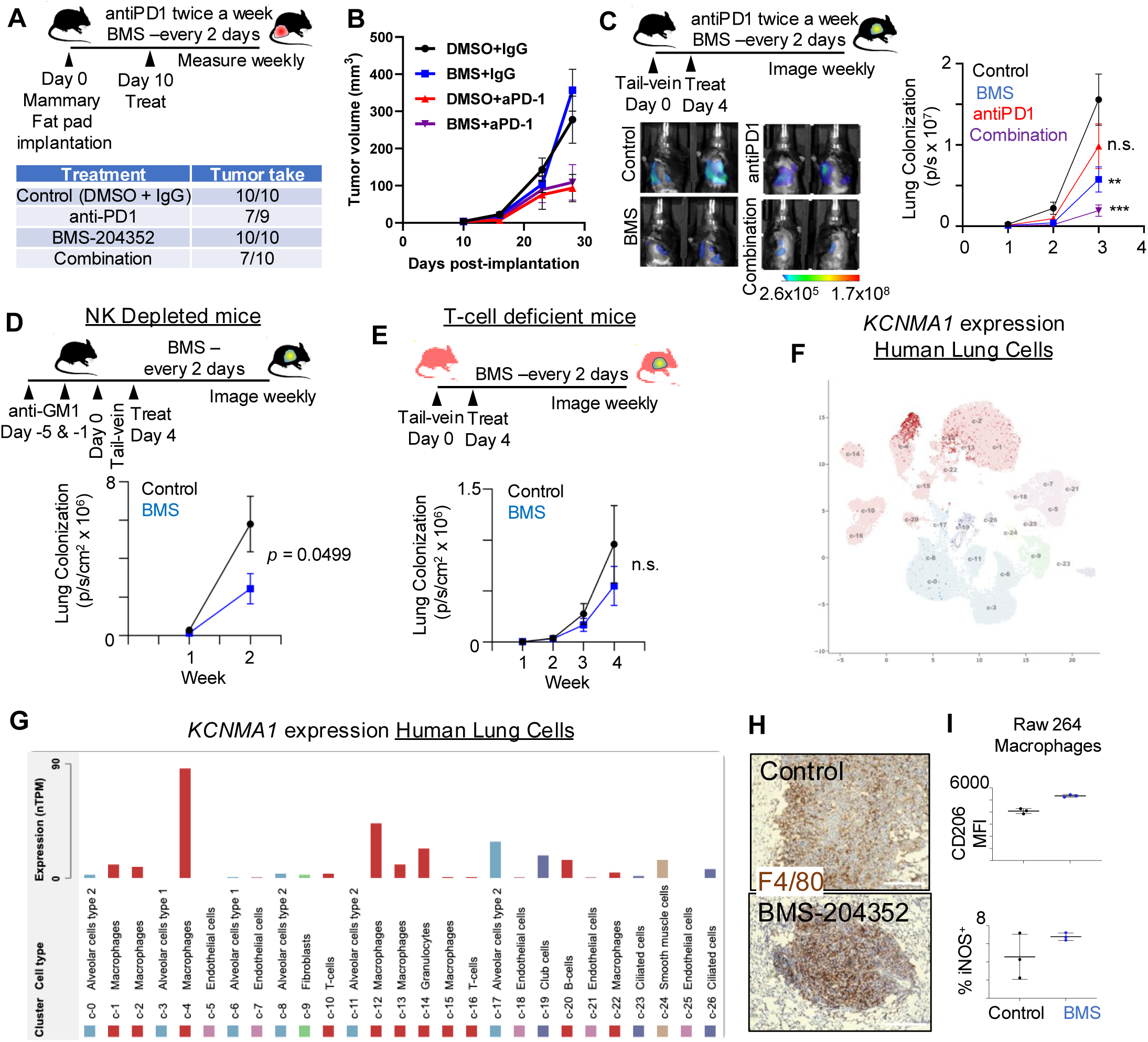
BK channel agonist treatment hinders metastatic colonization. (A) Orthotopic mammary fat pad tumor-take rates in mice injected with E0771 cells and treated with the indicated pharmacological agents. n= 9-10 female mice per treatment group (B) Orthotopic mammary fat pad tumor growth in mice listed in (A). Error bars, s.e.m. (C) Longitudinal analyses of metastatic burden in lungs of C57BL/6 mice over the course of 3 weeks by using bioluminescent imaging. N = 10 mice per treatment group. p/s: photons per second. P-values were derived from Mann-Whitney rank-sum test of two groups at a time. ns, not significant; \**P*<0.05; \*\**P*<0.01; \*\*\**P*<0.001. (D) Longitudinal analyses of metastatic burden in lungs of C57BL/6 mice pretreated with anti-GM1 antibodies to deplete NK cells. n = 10 female mice per group. p-value was derived from Mann-Whitney rank-sum test. (E) Longitudinal analyses of metastatic burden in lungs of athymic nude mice that lacks functional T-cells. n = 10 female mice per group. p-value was derived from Mann-Whitney rank-sum test. (F) UMAP representation of KCNMA1 expression in cells of the human lung accessed through the Human Protein Atlas database. Darker colors indicate higher degree of expression. Clusters c1-c26 correspond to the clusters c1-c26 in (G). (G) KCNMA1 expression in the clusters of cells represented in the UMAP as in (F). nTPM: transcripts per million. (H) Macrophage (F4/80) labeling of lungs from metastasis bearing mice that treated as in (C). Scale bar: 300µm. Images are representative of 8 mice in each group. (I) Macrophage polarization as judged by the surface expression of anti-tumor M1 marker iNOS and pro-tumor M2 marker CD206 in RAW 264 cell line. Representative of 2 independent experiments.

Next, we set out to determine the relative contribution of NK cells and T-cells to the anti-metastatic effects of BMS-204352. For this, we first used anti-GM1 antibody to deplete NK cells prior to injecting mice with E0771 cancer cells. BMS-204352 treatment in NK depleted mice could still hinder metastatic colonization, albeit not to the same degree as it did in fully immune competent mice (**Figure 6D**). BMS-204352 treatment in metastasis bearing athymic nude mice, which lack all T-cell populations, failed to significantly hinder metastatic colonization (**Figure 6E**). These data suggest that the functional presence of cytotoxic lymphocytes were necessary for BMS-204352 treatments to inhibit metastatic colonization, but they do not rule out contributions from other cell types in the tumor microenvironment (TME). Thus, we queried which cells of the tumor microenvironment expressed the molecular target of BMS-204352, KCNMA1 and its associated proteins KNCMB1 and JPH2 in cancer patients’ primary tumors and in healthy lung cells annotated in the Human Protein Atlas.^46–48^ KCNMA1 was detected in epithelial tumor cells, but its expression was most prominent in macrophages in primary breast tumors and in non-cancerous lungs (**Figures 6F**, **6G and S6K**). To determine whether BMS-204352 could alter macrophage recruitment, we used immunohistochemistry for the pan-macrophage marker, F4/80, in metastatic tissues. Metastatic foci in all conditions were heavily infiltrated by tumor associated macrophages (TAMs), and we did not note any obvious differences in the levels of TAM recruitment or morphology (**Figure 6H**). BMS-204352 treatment also did not significantly change pro- versus anti-tumorigenic properties of macrophages, as judged by its impact on the anti-tumor TAM marker iNOS or pro-tumor TAM marker CD206 in the murine Raw264 cell line model (**Figure 6I**). Together, these results suggest that cytotoxic lymphocytes, comprised of NK and T-lymphocytes were the cell type primarily responsible for pruning incipient metastatic colonies upon treatment with BMS-204352 *in vivo*.

### BK channel agonist treatment promotes cell stiffness under pathological ionic conditions

We sought to provide a deeper mechanistic explanation into why BMS-204352 reduced metastatic colonization. For this, we decided to model the TME with the appropriate ionic context. Standard cell culture media and human serum contain 3.5 – 5.5 mM extracellular potassium ([K^+^]_e_), but under pathological conditions serum [K^+^]_e_ can rise above 6.5mM. This elevated [K^+^]_e_ is called hyperkalemia, it is frequently observed in patients with tumor lysis syndrome (TLS) and it requires emergency medical intervention to address cardiac, muscle and neuronal dysfunction.^49,50^ Strikingly, electrolyte measurements of tumor interstitial fluid (TIF) and photoacoustic measurements of implanted tumors show that [K^+^]_e_ in the tumor microenvironment can reach up to 75 mM, an order of magnitude above clinically defined hyperkalemia.^51–53^ To determine whether there was similar elevation in [K^+^]_e_ in the lung metastatic niche, we loaded a polyacrylamide gel with a non-cell permeable K^+^ indicator (IPG-2 TMA Salt) and a cell permeable Hoechst dye (**Figure 7A**). Overlaying this gel to snap-frozen live sectioned lung tissues showed pockets of high [K^+^]_e_ areas consistent with the idea that elevated [K^+^]_e_ is a feature of the metastatic TME (**Figure 7B**). Thus, we measured the effects of BMS-204352 on cellular phenotypes both under physiological (5 mM) and pathological (45 mM) [K^+^]_e_. BMS-204352 treatment had modest and variable effects on cellular migration and proliferation under high and normal [K^+^]_e_ (**Figures S7A, S7B**). All treatments reduced the basal level of apoptosis, and this did not explain the anti-metastatic effect of BMS-204352 treatment (**Figure S7C**). Finally, we measured cancer cells’ sensitivity to lysis by cytotoxic T-lymphocytes. For this, we deployed the ovalbumin (OVA) model antigen approach where genetically engineered cytotoxic T-lymphocytes destroy cancer cells in a manner that is dependent on the concentration of the synthetic OVA peptide (SIINFEKL) presented by the cancer cells’ Major Histocompatibility Class I molecules.^54^ Activation of OVA specific T-lymphocyte was hindered by high [K^+^]_e_, as judged by Lamp1 positive cytotoxic granule presentation in their cell surface during co-culture with cancer cells and this was rescued by BMS-204352 treatment (**Figure 7C**). Importantly, cancer cell lysis by T-lymphocytes were also significantly slower under high [K^+^]_e_ and this was also rescued by BMS-204352 treatment (**Figure 7D**). Resensitization of cancer cells to cytotoxic T lymphocyte driven cell death correlated with cancer cell stiffness, where cancer cells cultured in high [K^+^]_e_ were significantly softer than cells cultured in regular media and this softening was rescued by BMS-204352 treatments (**Figure 7E-7F**).

**Fig. 7.**
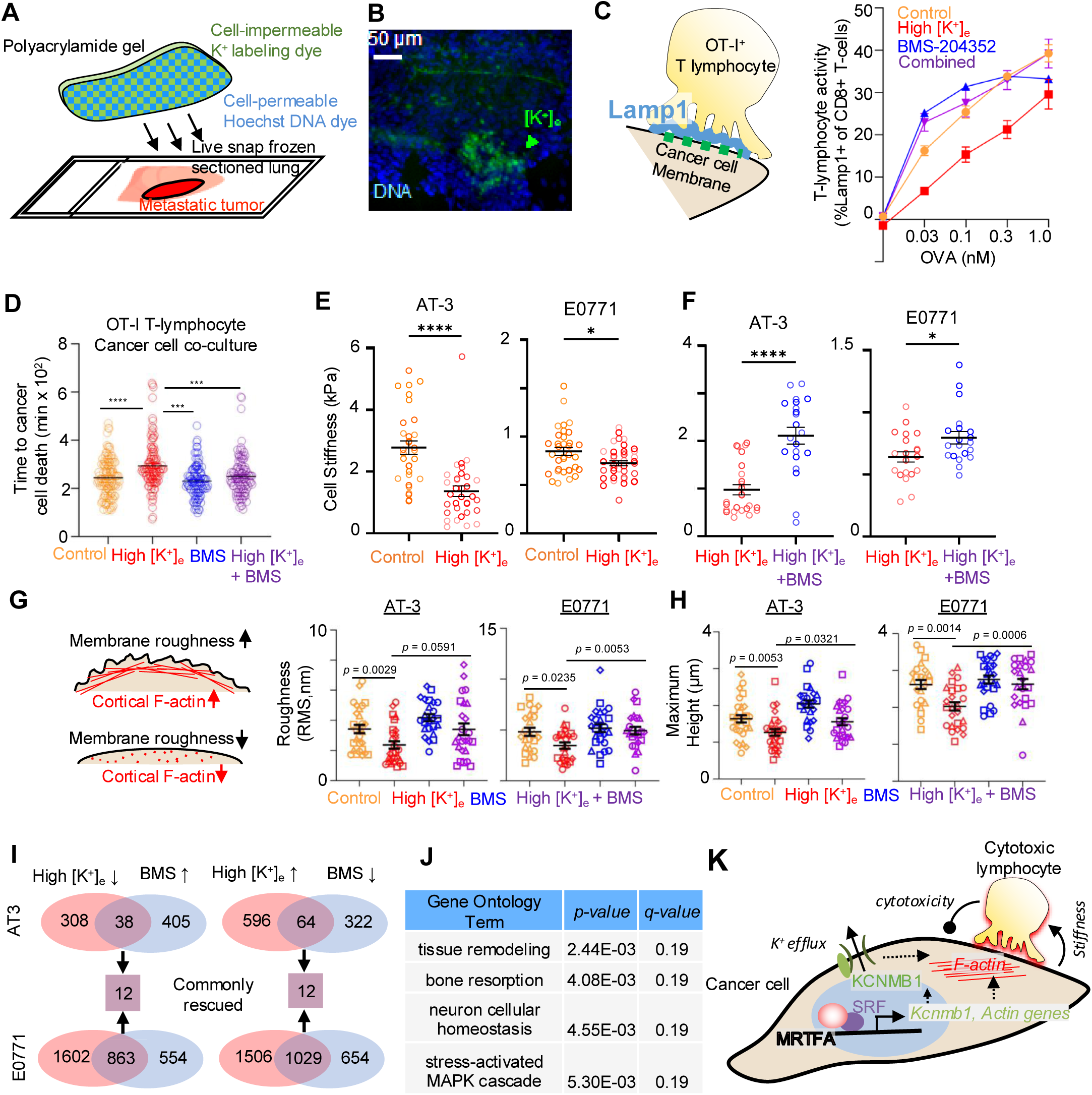
BK Channel activity promotes cellular stiffness in pathologically high [K^+^]_e_. (A) Schematic of the experimental approach to qualitatively assess relative amounts of extracellular potassium in the tumor microenvironment. (B) Representative image from a lung section showing accumulated K^+^ in a metastatic lesion. Green triangle points to accumulated [K^+^]_e_. Scale: 50µm. (C) Schematic showing elevated Lamp1+ granules at the immune synapse during antigen mediated cancer cell targeting by OT-I+ CD8+ Cytotoxic T lymphocytes. Graph on the right shows OVA dependent increase in Lamp1+ positivity as a measure of T-lymphocyte granulation under different conditions. Results are representative of 3 independent experiments. (D) Measurements of time to death of a cancer cell after it has been contacted 1:1 by a CD8^+^ T lymphocyte as determined by PI accumulation in the cancer cell. Data are compiled from 4 independent experiments with. P-values are calculated by Kruskal-Wallis test with multiple comparisons. Mean values are shown. ***:P<0.001, ****:P<0.0001. Control = 119 cells, high [K^+^]_e_ = 117 cells, BMS = 107 cells, high [K^+^]_e_+BMS = 109 cells. (E) AFM stiffness measurements of representative cancer cells treated with high [K^+^]_e_ (45 mM KCl) compared to control. Results are from 2 combined experiments. AT-3: Control = 30 cells, high [K^+^]_e_ = 33 cells; E0771: Control = 34 cells, high [K^+^]_e_ = 37 cells. Mann-Whitney test. All error bars are mean±s.e.m. ns, not significant; *P<0.05; **P<0.01; ***P<0.001. (F) AFM stiffness measurements of representative cancer cells treated with high [K^+^]_e_ and with/without 10 µM BMS-204352. Results are from 2 combined experiments. AT-3: high [K^+^]_e_ =25 cells, high [K^+^]_e_+BMS = 21 cells; E0771: high [K^+^]_e_ = 20 cells, high [K^+^]_e_+BMS = 20 cells. Mann-Whitney test. All error bars are mean±s.e.m. ns, not significant; *P<0.05; **P<0.01; ***P<0.001. (G) Schematic depiction of how cortical F-actin influences surface roughness and the surface roughness measurements performed by AFM. Data is combined from 4 independent experiments. AT-3: Control = 28 cells, high [K^+^]_e_ = 28 cells, BMS = 24 cells, high [K^+^]_e_+BMS = 24 cells, E0771: Control = 23 cells, high [K^+^]_e_ = 25 cells, BMS = 24 cells, high [K^+^]_e_+BMS = 23 cells. P-values calculated by Mann-Whitney rank sum test. Mean+/-s.e.m are shown. RMS: root-mean square. (H) Cell height of cells used in (G) measured by AFM. (I) Venn diagrams showing the number of differentially regulated genes in high [K^+^]_e_. The intersections represent the genes “rescued” which means the BMS-204352 treatment caused differential expression in the opposite direction. “Commonly rescued” genes are conserved across two different cell types. (J) Gene ontology analyses of “Commonly rescued” genes in (F). See Table S6 for a full list of GO terms that are enriched. (K) Working model: MRTFA/SRF upregulates the expression of F-actin bundling proteins and BK channel subunit, KCNMB1, for promoting cellular stiffness. Elevated cancer cell stiffness triggers mechanosurveillance by cytotoxic lymphocytes for inhibiting metastatic colonization.

Finally, we set out to determine how BMS-204352 treatment elevated cellular stiffness. Unlike the genetic perturbations, there was no difference in F-actin stress fiber alignment in various conditions (**Figure S7D**). We reasoned that anisotropy measurements may not capture stiffness changes in cortical actin associated with plasma membrane. Thus, we measured plasma membrane roughness, which is an established secondary measure of cortical actin assembly.^55,56^ We found that high [K^+^]_e_ significantly smoothened cell membranes and lowered cell height and this was rescued by BMS-204352 treatment, suggesting that BMS-204352 mediated cell stiffness involves direct regulation of cortical actin assembly (**Figure 7G and 7H**). We also considered two alternative explanations for the cell stiffness phenotypes: BMS-204352’s impact on transcriptional regulation and on mitochondrial organization. To investigate how BMS-204352 could regulate cell stiffness via transcriptional regulation, we performed bulk RNA-sequencing under various conditions. A total of 1006 and 5000 genes were differentially regulated by high [K^+^]_e_ in AT-3 and E0771 cell lines, respectively. Of these, the expression of 24 genes were commonly rescued in both cell lines, but there were no biological processes that involved cytoskeletal elements and that showed a statistically significant enrichment with a false discovery rate of less than 10% (**Figures 7I-J and Table S5**). Mitochondrial fission and fusion can affect overall stiffness of the cell and BMS-204352 may regulate mitochondrial BK channels.^57,58^ To test this, we measured mitochondrial length and aspect ratio under different treatments, but they were not significantly affected either (**Figure S7E**). Together, these data show that BK channel activation regulates cancer cell stiffness through regulation of the F-actin cytoskeleton (**Figure 7K**).

## DISCUSSION

Here, we used a variety of gain-of-function and loss-of-function approaches to reveal a previously unrecognized role of MRTFA/SRF in regulating Ca^2+^ and K^+^ handling genes’ expression and ionic content in cancer cells. This was surprising finding because SRF is typically thought to be a downstream effector of Ca^2+^ signaling and Ca^2+^-induced cellular contractility. For example, pharmacological and genetic inhibition of the ion channel Transient Receptor Potential Cation Channel Subfamily M Member 7 (TRPM7) in liver cancer cells prevents RhoA-mediated MRTFA/SRF nuclear localization and transcriptional activity.^59^ Similarly, VGCC activity stimulates SRF binding to its target CArG boxes in the genome for vascular smooth muscle cell contraction.^60^ Conversely, our results indicate several ion channel genes are regulated by MRTFA/SRF. While we focused on Ca^2+^ and K^+^ ions, it is possible that MRTFA/SRF regulates other monoatomic ion transport such as sodium because we also found sodium channel genes, such as *Scn5a*, to be upregulated by MRTFA^WT^ overexpression. This is consistent with the high abundance of SRF target CArG boxes in the promoters of genes that define smooth muscle cell identity to properly perform EC coupling for their physiological function.^61–63^

We found that a major contributor to MRTFA/SRF-mediated cortical stiffening in cancer cells was KCNMB1, the auxiliary subunit of the BK channels that promotes the open-state of BK channels by sensitizing them to elevations in local [Ca^2+^]_i_. MRTFA/SRF also regulated the expression of JPH2, which promotes K^+^ efflux by bridging CICR channels in the ER/SR to KCNMB1 and the BK channels.

Interestingly, our findings in cancer cells contradict findings in electrically excitable cells, where the loss of these proteins is expected to promote stiffening either through continued Ca^2+^ leakage from ER/SR or through continued opening of VGCC channels and thereby promoting contractility. Based on our RNA sequencing data, we reason that stiffness modulation is different between cancer cells and electrically excitable cells because cancer cells do not express VGCCs at sufficient levels. Consistent with this idea, we also did not find changes in [Ca^2+^]_i_ when KCNMB1^E65K^ was overexpressed. Together, these results suggest that KCNMB1 promotes cellular stiffness and F-actin polymerization by increasing K^+^ efflux without a compensatory increase in [Ca^2+^]_i_, but the precise mechanism of F-actin regulation remains unknown. One possibility is that K^+^ efflux promotes hyperpolarization of the plasma membrane, which changes the electrical charge of the lipid bilayer and thereby regulating the electrical attraction and repulsion of the filaments.^64,65^

Ionic regulation of cancer progression is an emerging area of investigation. Particularly, potassium homeostasis has drawn recent attention because tumor interstitial fluid contains severely elevated [K^+^]_e_.^51–53^ This is a remarkable finding because even the smallest changes in serum [K^+^] outside the normal range (3.5 – 5.5 mM) requires medical intervention and small changes in intracellular potassium has functional consequences in cell biological processes.^49,66^ Elevated [K^+^]_e_ has been proposed to inhibit cytotoxicity by interfering with AKT signaling in T-lymphocytes, and our results revealed a previously unrecognized biophysical dimension to lymphocyte dysfunction: Elevated [K^+^]_e_ reduces cortical stiffness, which is critical for effective perforin granule insertion and mechanosurveillance: the mechanochemical activation of lymphocytes at the immune synapse.^6–14^ The nature of cortical stiffening dictates the mechanism of cytotoxic action. For example, hypertonic challenge and the subsequent loss of membrane tension is sufficient to inhibit lymphocytes’ ability to insert perforin and granzyme into the target membrane.^11^ Similarly, cholesterol depletion, which stiffens the cell cortex at the immune synapse facilities F-actin orientation for better effective force exertion, but these modes of mechanics do not seem to effect mechanotransduction in lymphocytes.^6^ Conversely, regulation of the underlying actin cytoskeleton does affect lymphocyte mechanotransduction and involves mobilization of Lamp1^+^ cytotoxic granules to the cell surface.^9^ In the case of high [K^+^]_e_ and BMS-204352 treatments, we find that these treatments affect Lamp1^+^ granules to the cell surface and this correlates with heightened T-lymphocyte mediated killing of co-cultured cancer cells. Future studies will investigate how elevated [K^+^]_e_ and the rescue of cancer cell stiffness through pharmacological perturbation, such as BMS-204352, affect other aspects of mechanosurveillance in lymphocytes such as force exertion at the immune synapse and long-term transcriptional affects in T-lymphocytes.

An exciting outcome of our work is the potential pre-clinical and clinical application of ion channel modulators to alter cancer cell stiffness and metastasis. Existing therapies target genetic and biochemical abnormalities in cancer cells, while overlooking the biophysical ones. This is because cancer cell specific genetic and biochemical abnormalities provide a broad therapeutic window to target these processes, whereas cancer cell specific mediators of biophysical characteristics, such as stiffness, are not known. Here we described KCNMB1 mediated K^+^ efflux as a cancer cell specific modulator of cell stiffness, since electrically excitable, non-transformed pericytes behaved the opposite way under the same genetic perturbations. We also found that the prolonged use (3 weeks) of BK channel activator, BMS-204352, is safe from a cardiac health perspective in our pre-clinical model.

Importantly, BMS-204352 has demonstrated clinical safety when used in human trials to treat stroke patients, which supports the idea that BMS-204352 could be repurposed for uniquely promoting stiffness of cancer cells and thereby limit metastatic colonization in patients.^43^ Agonism and antagonism of various potassium channels including BK channels have been extensively studied and found to be tumor-promoting or tumor-inhibiting in different *in vitro* and immune deficient xenograft models.^67–74^ Our studies in immune competent mice underscore the importance of immune context in evaluation of potassium channel perturbations for cancer treatment and highlight the biophysical dimension of bioelectricity, immunity and metastasis.

## LIMITATIONS OF THE STUDY

Even though our working model suggests that KCNMB1 protein expression would be negatively selected against in metastatic tissues, we could not test this because KCNMB1 antibodies did not reliably detect mouse KCNMB1, as previously reported.^75^ In the present study, we do not provide data for how [K^+^]_e_ levels, lymphocyte activation and cellular stiffness are related *in situ*. This is because the ionic microenvironment and lymphocyte activation through biophysical means require intact tissues and cell-cell contact, which would be disrupted during traditional preparation of lymphocytes for activity measurements in *ex vivo* flow cytometry or *in situ* histology approaches. Thus, further support for our model requires new technologies to be developed to study the biophysical impact of the ionic microenvironment on immune function.

## METHODS

### Recombinant DNA

shRNA constructs for both mouse and human were purchased from Sigma in the pLKO.1 system (Mouse: KCNMB1 #TRCN0000068366, SRF #TRCN0000054593, JPH2 #TRCN0000423179; Human: KCNMB1 #TRCN0000043794, SRF # TRCN0000010838). shMRTFA/B was previously described ^9^. pLKO.1 was used as control for all knockdown experiments. The MRTFA^Y238A^ and KCNMB1^E65K^ mutants were created by single point mutations in *MRTFA* (Addgene #19846) and *Kcnmb1* (Addgene #113565) and were verified by Sanger sequencing. To generate overexpression of the wild type and mutated constructs, coding regions of the genes were PCR amplified and subcloned into retroviral vector pRetroX-Tight-Hygro (Takara Bio #631034). pRetroX-Tight-Hygro was used as control for overexpression studies. For doxycycline-inducible cell line generation, pLVX-TetON-Advanced (rtTA) (Takara Bio, #632162) was used. To over express YAP (Addgene #33091) and its active mutant YAP^5SA^ (Addgene #33093) were cloned into pLVX-puro (Takara Bio Catalog No. #632164).

### Cell culture

AT-3 cells were cultured in DMEM (Fisher Scientific # MT10017CV) containing 10% (vol/vol) FBS (Thermo Fisher # 26140079), 1 mM sodium pyruvate (Fisher Scientific #11-360-070), 2 mM non-essential amino acids (Thermo Fisher #11140050), 15 mM HEPES (Fisher Scientific #15-630-080), and 100 µM B-mercaptoethanol (Sigma #ES-007-E). E0771 cells were cultured in RPMI (Fisher Scientific #MT15040CV) containing 10% (vol/vol) FBS, 15 mM HEPES, 100 U/mL penicillin and 100 µg/mL streptomycin (Fisher Scientific #10-378-016). SKBR3 cells were cultured in McCoy’s 5A media (ATCC #30-2007) supplemented with 10% (vol/vol) FBS. BT-474 cells were cultured in DMEM containing 10% (vol/vol) FBS. Pericytes were cultured in Pericyte Medium (ScienCell #1201). All cells were incubated at 37°C with 5% CO_2_. Drug selection for genetically engineered cells was achieved by culturing cells with 1000 µg/mL G418 (Sigma #108321-42-2), 500 µg/mL hygromycin (Sigma #10843555001) and 2 µg/mL puromycin (Sigma #58-58-2). Overexpression was induced by treating cells with 2 µg/mL doxycycline (Fisher Scientific #10592-13-9) 48 hr prior to use in experiments.

### RNA sequencing

1.5 x 10^5^ AT-3 or E0771 Control (pRetroX), MRTFA^WT^, and MRTFA^Y238A^ were seeded in a 6-well plate for 48 hr with doxycycline treatment to induce overexpression. Cells were then collected and processed with the RNeasy mRNA extraction kit (Qiagen #74106) according to the manufacturer’s instructions with the inclusion of the DNase treatment step. RNA samples were then submitted to the UIC Genome Research Core where they were checked for purity using NanoDrop™ One Spectrophotometer (Thermo Scientific) and analyzed for integrity using Agilent 4200 TapeStation and RNA ScreenTape. Relative levels of remaining DNA were checked by dual RNA/DNA measurements using Qubit fluorometer (Invitrogen). DNA amounts did not exceed 10% of the total amount of NA. Illumina sequencing libraries were prepared using 250 ng of total RNA per sample. Library preparation was carried out with the NuGen Universal Plus mRNA w/ NuQuant (Tecan Genomics former NuGen), as per the manufacturer’s protocol rev V6. In brief, the library construction steps consist of poly(A) RNA selection, RNA fragmentation and double-stranded cDNA generation using a mixture of random and oligo(dT)priming. The library is then constructed by end repairing the cDNA to generate blunt ends, ligation of UDI adaptors, strand selection, and PCR amplification. The required number of PCR cycles was determined to be 10 by qPCR on a small aliquot of the un-amplified libraries. Final amplified libraries were quantified with the Qubit 1X dsDNA HS Assay Kit (Invitrogen), and fragment size distribution was confirmed to be approximately 270 bp using the D5000 ScreenTape assay (Agilent). The concentration of the final library pool was confirmed by qPCR and subjected to a test sequencing run on a MiniSeq instrument (Illumina) in order to check sequencing efficiencies and adjust accordingly proportions of individual libraries. The adjusted library pool was re-quantified by qPCR using Library Quantification Kit (Kapa Biosystems) and sequenced on the NovaSeq 6000 instrument (Illumina), S4 flowcell, 2×150 bp reads. Fastq files were quality checked using FastQC (https://www.bioinformatics.babraham.ac.uk/projects/fastqc/) prior to analysis. Reads were aligned using STAR (version 2.7.6a) ^76^ against the GRCm38 (mm10) genome provided by Ensembl. Count tables were generated using featureCounts ^77^ (Subread release 2.0.1). Data are available through GEO Datasets GSE273414 and GSE285078. Differential expression analysis was performed using Deseq2^78^. Gene set enrichment analysis was performed with clusterProfiler (10.1016/j.xinn.2021.100141) against the GO Biological Processes database (10.1093/nar/gkm883). The background gene set was determined by genes detected above an average count of 10 reads in the respective RNA sequencing data. Enrichment scores with a *q-value* (false discovery rate) *<0.05* were considered statistically significant.

### RT-PCR and ChIP-PCR

1.5 x 10^5^ cells with overexpression constructs were seeded in a 6-well plate for 48 hours then collected and processed with the RNeasy mRNA extraction kit (Qiagen #74106) according to the manufacturer’s instructions. The resulting RNA was then reverse transcribed into cDNA according to the High-Capacity cDNA Reverse Transcriptase Kit from Applied Biosystems (#4368814). Applied Biosystem’s TaqMan Fast Advanced Master Mix for qPCR (#4444556) was used according to the manufacturer’s instructions to prepare the samples for RT-PCR. TaqMan probes used were all purchased from Thermo Scientific: *Gapdh* (Mm99999915_g1), *Acta2* (Mm00725412_s1), *Actg2* (Mm00656102_m1), *Tagln* (Mm00441661_g1), *Myh11*(Mm00443013_m1)*, Mrtfa* (Mm00461840_m1), *Mrtfb* (Mm00463877_m1), *Srf* (Mm00491032_m1), *Kcnmb1* (Mm00466621_m1), *Ccn2* (Mm01192933_g1), *GAPDH* (Hs99999905_m1), *MRTFA* (Hs01090249_g1), *MRTFB* (Hs00401867_m1), *SRF* (Hs00182371_m1), *KCNMB1* (Hs00188073_m1). Each sample was run with 3 technical replicates on a 384-well plate on the ViiA 7 Real-Time PCR System (Thermo Fisher) housed by UIC’s Genome Research Core and the data was processed using the system’s corresponding QuantStudio (Thermo Fisher) software. Chromatic immunoprecipitations (ChIP) were performed with SimpleChIP® Plus Enzymatic Chromatin IP Kit (Magnetic Beads) from Cell Signaling Technology (Cat#. 9005) as manufacturer’s instructions. Histone H3 (D2B12) XP Rabbit mAb (Cat#. 4620, Cell Signaling Technology), Normal Rabbit IgG (Cat#. 2729, Cell Signaling Technology), SRF (D71A9) XP Rabbit mAb (#5147, Cell Signaling Technology), and MKL1/MRTF-A (E2V2I) Rabbit mAb (Cat#. 97109, Cell signaling Technology) were used for immunoprecipitation. PCR was performed using KOD Hot Start DNA Polymerase (Cat#. 71316, Novagen) according to the manufacturer’s protocol with primers targeting the KCNMB1 gene; Forward: 5’ - AGACTCTGGGAAATGCAGGG – 3’, Reveres: 5’ - GGAAGCAGTCTCAGTCCCAT – 3’. Immunoprecipitated DNAs and 2% input DNA were used as templates.

### Immunoblotting

250,000 cells were seeded in 6-well dishes for 48 hours, washed with ice cold PBS and lysed with RIPA lysis buffer (Cell Signaling #9806), containing proteinase and phosphatase inhibitors. Cell lysates were transferred to 1.5 mL tubes on ice for 10 min, sonicated on ice for 30 seconds, lysed on ice for 30 min and spun for 5 min at 13,000 RPM. Lysates were transferred to new tubes and 50 µL of Laemmli loading buffer (Bio-Rad Laboratories #1610747) was added to each sample. Samples were boiled at 90°C for 7 min, then loaded and run on pre-cast 4-12% gels (Invitrogen #NP0336BOX) in MOPS buffer. Membrane transfer was performed using the iBlot Transfer System (Thermo Scientific) and the resulting membranes were blocked for 1 hour in 5% milk in TBS-T at room temperature. Primary antibodies were also diluted in 5% milk in TBS-T and added to the membranes for overnight rocking at 4°C. The following day, membranes were washed with TBS-T for 5 min, 3 times. Secondary antibodies were diluted the same as the primaries and added to the membranes for 1 hour. Membranes were dried on blotting paper and imaged using the BioRad ChemiDoc Imager. Antibodies used: GAPDH 1:2000 (Cell Signaling #2118S), YAP 1:1000 (Cell Signaling #14074S), pSMAD2/3 1:1000 (Cell Signaling #8828S), SMAD2 1:1000 (Cell Signaling #5339S), MRTFA 1:500 (Cell Signaling #97109S), Goat anti-Mouse IgG (H+L) Cross-Adsorbed Secondary Antibody, DyLight™ 800, Invitrogen™ 1:10,000 (Thermo Scientific #SA510176), Goat anti-Rabbit IgG (H+L) Cross-Adsorbed Secondary Antibody, DyLight™ 800, Invitrogen™ 1:10,000 (Thermo Scientific SA510036).

### Atomic force microscopy

Cells were plated on glass-bottom Fluoro dishes (World Precision Instruments), cultured for 48 hours in their appropriate cell culture media and subjected to atomic force microscopy measurements. Between 4-8 cells per condition per day were measured on the Asylum MFP-3D SPM. We use colloidal probes, (Novascan, PT.GS and Bruker, MLCT-SPH-5UM) with sphere radius of 2.5 μm and nominal spring constant of 0.06 N/m and 0.08N/m. The exact spring constant of the cantilever was determined by thermal tune approach after establishing the deflection sensitivity prior to each experiment. Force-indentation maps of 60×60 μm2 (18×18 points) in areas containing cells was generated at indenting velocity of 2 μm/s and targeted external force stimulation of 1 nN. Cellular stiffness (Young’s modulus) was calculated by applying the Hertzian model to the force indentation curves yielded by the maps.

For the roughness measurements, topography maps were acquired with the same probe, A 40×40 μm^2^ (32×192 points) area was scanned at a scan rate between 0.1 and 0.2 Hz. To select a representative area for analysis, the maximum point of the cell was found, and the corresponding row, along with the rows immediately above and below, were selected to analyze. The analysis software Gwyddion was used for data evaluation, to measure the mean square roughness RMS and the mean roughness of the cellular membrane.^79^ RMS represents the height irregularities with the mean and was computed 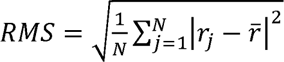 across all points.

### Immunofluorescence and microscopy

5,000 cells were seeded per well in an 8-well glass chamber slide and cultured for 48 hours, fixed with 4% paraformaldehyde and stained for either F-actin using Acti-Stain 555 Phalloidin (Cytoskeleton #PHDH1-A) according to the manufacturer’s instructions or with MRTFA antibody 1:500 (Cell Signaling #97109S) overnight, following 1 hr of blocking with 10% normal goat serum, 2% BSA, and 1% Triton-X. After washing 3 times with 1% Triton-X solution, the secondary antibody (Goat anti-Mouse IgG (H+L) Secondary Antibody, Alexa Fluor® 555 conjugate) for MRTFA was added 1:1000 for 1.5 hours. Finally, the slide was stained with Hoescht nuclear stain for 10 minutes and was mounted with Immu-Mount mounting media (Fisher Scientific #9990412) and glass coverslip. Confocal images were taken with the 63x objective lens on the Zeiss LSM 880 Confocal. A minimum of 30 images of at least one well were taken per cell line, per experiment. For staining with propidium iodide to determine relative cell death induced by EDTA or BAPTA-AM treatments, propidium iodide was used at a final concentration of 6 µM and brightfield/RFP channel images were taken using a 10x objective lens.

### Drug treatments and doses

Cells were pre-treated with the following reagents for the indicated times and the indicated doses: SB-505124 (2.5 µM; 24 hrs) (ApexBio #A3799); Verteporfin (10 µM; 24 hrs) (ApexBio #A8327); EDTA (1 mM; 20 min) (RPI #E14000-100.0); BAPTA-AM (10 µM; 1 hour) (Thermo Scientific #B6769); KCl (40 mM; 24 hours); BMS-204352 (10 µM; 24 hours) (A2Z Chemicals/STARK Chemicals #BMS204352).

### Fluorescent labeling assays

For both ion labeling assays, 10,000 cells were seeded per well in a black walled glass bottom 24-well plate for 48 hour prior to the experiment. Before imaging for the calcium labeling assay, cell media was aspirated and replaced with 1 mL PBS containing 2 µM Fluo-4 AM (Thermo Fisher #F14201), 5 µM Fura Red (Thermo Fisher #F3021), and 1x PowerLoad (Thermo Fisher #P10020). The cells were incubated at 37°C for 30 min, upon which the solution was aspirated and phenol red-free RPMI supplemented with 10% FBS was added to the cells for imaging. Using the 20x objective lens on the Zeiss LSM 880 Confocal Microscope, one 4×4 stitched image was taken per well, constructed from a Z-stack of 10 slices spanning a total of 9 µm. These images were captured using a laser excitation at 488 nm and emissions at 539 +/- 40 nm (green) and 682 +/- 70 nm (red) as per the preset values for Fluo-4AM and Ca^2+^ Free Fura-Red in the Zeiss ZEN software. Potassium labeling with 500 - 2500 ng/mL IPG-1AM was conducted per manufacturer protocol using probenecid (ION Biosciences #3041F). After the 1 hour incubation of the dye loaded plate at 37°C, the dye solution was removed and replaced with dye-free RPMI media with 10% FBS for imaging using 525 nm excitation and 545 nm emission settings. For extracellular K^+^ staining a 250µL polyacrylamide gel was polymerized by mixing 38.6µL of 30% ProtoGel, 198.2µL ddH_2_O, 1µL Hoechst dye, 6.125µL IPG-2 TMA^+^ Salt (2 µg/µL dissolved in water), 1µL of TEMED and 5µL of 10% APS. The mixture was put in a glass mold with approximately 0.34 mm thickness and flattened out with a coverslip placed on top. The resulting gel was place over 10µm cryosectioned metastasis bearing mouse lung tissue that was previously snap frozen without perfusion. The gel and tissue were immediately imaged on an EVOS M5000 upright bright field microscope. Mitochondrial staining was performed by using MitoTracker Green FM (Thermo Scientific, M7514) per manufacturer’s instructions.

### Electrophysiology

Manual whole-cell patch clamp analyses were performed on AT3 cells expressing KCNMB1^WT^ or KCNMB1^E65K^. Solutions and patch-clamp protocol were previously described.^80^ Specifically, extracellular solution contained 90 mM NaCl, 60 mM KCl, 10 mM HEPES, 1.5 mM CaCl2, 1 mM MgCl2, 1 mM EGTA, pH 7.3. Pipette solution contained 145 mM KCl, 10 mM HEPES, 1 mM MgCl2, 4 mM ATP, 1 mM EGTA, pH 7.3. Intracellular solution was supplemented with 50 µM Ca2+ Maximum inward current was assessed by 500 ms linear voltage ramps from 0 mV to 80 mV. The holding potential was − 30 mV and pipette and whole-cell capacitance was automatically compensated.

### Cell viability and death assays

750 cells were seeded in 96-well plates in triplicate. Cells were treated with drugs 24 hours after seeding and kept in culture for up to 2 - 4 days. A control plate was analyzed on Day 1 prior to drug treatments and these values were used for normalization. The CellTiter-Glo Luminescent Cell Viability Assay (Promega #G7572) was used according to the manufacturer’s instructions to measure cell viability on the SpectraMax i3x Multi-Mode Plate Reader (Molecular Devices). Cell death was determined by caspase 3/7 cleavage using the Caspase-Glo 3/7 Assay (Promega #G8091) according to the manufacturer’s instructions 24 hours post-treatment with the inhibitors and by using the SpectraMax plate reader.

### Lymphocyte-cancer cell co-culture experiments

Mouse NK cells were freshly isolated from C57BL/6 splenocytes by negative selection using an NK cell isolation kit (STEMCELL Technologies #19855) and incubated overnight in 1000 U/ml IL-2. Mouse NK media: RPMI containing 10% FBS, PenStrep (1x)/L-Glutamine, and B-mercaptoethanol (1x or 0.1mM) 2,500 target AT-3 cells were seed in a black 24-well dish with a thin glass bottom in 1 mL of media the day prior to the start of the co-culture timelapse. Mouse NK cells were also freshly isolated on this day and cultured on their own. On the day of the co-culture experiment, NK cells were collected and stained with CellTrace Violet stain (Thermo Scientific #C34557) according to the manufacturer’s instructions. After staining, 25,000 NK cells were added to each AT-3-containing well along with 1000 U/mL IL-2 and 6 µM propidium iodide to mark cell death. The Zeiss LSM 880 Confocal Microscope was used for timelapse imaging overnight (18 hrs), complete with an incubation chamber which kept cells at 37°C and 5% CO_2_ during the course of the experiment. Images were taken every 6 min to produce a timelapse video. Quantitation of NK-mediated cell deaths was achieved by manual observation of 1:1 NK contact with a cancer target cell, then counting the number of frames it took for the target cell to shrivel up as a result of membrane blebbing and ultimately accumulate propidium iodide at which point the cell was registered as dead. For OT-I CD8^+^ T-lymphocyte cancer cell co-culture experiments, CTLs were obtained from splenocytes of OT-I mice and cultured with synthetic antigen OVA and IL-2 as previously described.^9^ OT-I CTLs were used between Day 5-7 after isolation. Cancer cells were incubated with the indicated treatments 24 hours prior to loading with 0.1 nM OVA and co-culturing with OT-I CTLs. Quantitation of OT-I CTL and NK-mediated cell death was conducted in a blinded manner. For OT-I CTL and cancer cell co-culture experiments, CTL activity was measured by adding exogenous Lamp1 antibody for 90 minutes to the cell culture media to exclusively label surface Lamp1 expression.

### Tumor growth and metastasis experiments

For orthotopic implantation of tumor cells, 50,000 E0771-TGL cells were resuspended in a 50/50 mixture of PBS and growth factor reduced phenol red free Matrigel in 50 µLs of volume and implanted into the 4^th^ mammary fat pad of 4 – 6-week-old female C57BL/6J (Jackson Labs 000664) mice by exposing the mammary fat pad through an incision into the skin as per approved UIC Animal Care Committee protocols. For experimental lung colonization, 300,000 E0771-TGL cells were injected into the tail vein of 4 – 6-week-old female C57BL/6 mice or 4 – 6-week-old Hsd:Athymic Nude-Foxn1nu mice (Envigo-Inotiv Stock #069). Only female mice were used for all experiments because 99% of all breast cancers happen in females. Orthotopic tumor growth was measured weekly by calipers and lung metastasis burden was quantified weekly using retro-orbital D-luciferin (150 mg kg^−1^) injection followed by imaging via the LagoX Imager (Spectral Instruments Imaging). 200 µg anti PD-1 (BioXcell #BP0146) or mouse IgG control (BioXcell #BP0091) antibodies were injected intraperitoneally per mouse, bi-weekly. NK depletion was achieved by injecting 33 µg anti-asialo-GM1 antibody (Wako Chemicals) 6 days and 1 day before tumor cell injection and every 5 days thereafter. 3 mg/kg BMS-204352 (A2Z Chemicals/STARK Chemicals #BMS204352) or DMSO control solutions were also injected intraperitoneally every other day, and prepared fresh each time using the formulation 5% Tween 80, 40% PEG 400, 5% DMSO.

### Immunohistochemistry of mouse tissues

Paraffin blocks with mouse tumors were sectioned at five micrometers. A subset of slides was used for immunohistochemistry with F4/80 antibody (1:250, 70076, Cell Signaling). Staining was performed with BOND Polymer Refine Detection Kit (Leica, DS9800) on BOND RX automated stainer (Leica Biosystems) according to the preset protocol. After deparaffinization, sections were subjected to heat-based antigen retrieval with BOND Epitope retrieval buffer 1 (pH 6.0, Leica Biosystems, AR9961) for 20 min at 99°C. Endogenous peroxidase activity and non-specific binding was blocked by sequentially treating samples with peroxidase block (BOND Polymer Refine Detection Kit) and protein block (Background Sniper, Biocare Medical, BS966) for 15 min at room temperature. Sections were then incubated with F4/80 antibody for 30 min. The signal was detected with anti-rabbit-Poly-HRP and DAB from BOND Polymer Refine Kit by incubating sections for 15 min and 10 min at room temperature correspondingly, and hematoxylin was used as a counterstain. All slides were dehydrated on Autostainer XL and mounted with Micromount media (Leica Biosystems)

### Mouse echocardiography

Echocardiography was performed on control and drug treated mice 4 days and 2.5 weeks after tail-vein injection of cancer cells using the Vivo 2100 with the MS550D transducer. BPM was maintained between 400 - 600 BPM and mice were maintained on 1% isoflurane throughout the procedure. Echocardiography was performed on female mice, and the following images were taken: Left ventricular systolic function: long axis & short axis left ventricle images (m-mode) were taken. Left ventricular diastolic function: Apical 4 chamber (color doppler and tissue doppler). Right ventricular function: Right ventricular free wall and pulmonary artery blood flow. Data using analyzed with Graphpad Prism and statistics performed included: two-way ANOVA with post-hoc TukeyHSD.

## QUANTIFICATION AND STATISTICAL ANALYSIS

### Statistics

Each experiment was repeated independently at least twice, and statistics were performed on each representative experiment or combined results as stated in the figure legend. When data were combined from multiple experiments, each experiment was distinguished by different color shades or shapes. Data were tested for binomial distribution before statistical analyses. GraphPad Prism was used to perform all statistical tests, including: Kruskal-Wallace with multiple comparisons, Mann-Whitney, two-tailed ANOVA, and paired and unpaired two-tailed t tests. All error bars denote SEM unless otherwise noted. Sample sizes were not determined by statistical methods prior to experiments.

### TCGA data analysis

TCGA patient data for analyzing the expression of MRTFA target genes, such as KCNMB1 and KCNV2, in MRTFA^high/low^ and SRF^high/low^ patients were accessed by using cBioportal (Memorial Sloan Kettering Cancer Center). To determine the enrichment of calcium gene signatures TCGA data for invasive breast cancer PanCancer Atlas downloaded through the Genome Data Commons (Broad Institute, Massachusetts). Gene Signature Enrichment Analysis (GSEA) v4.3.3 was used to determine the enrichment of gene sets that included the key words “monoatomic”, “transport”, “channel” using default settings inclusive of all gene sets regardless of size.

### Image analysis

F-actin alignment, or anisotropy, in phalloidin-stained cells was quantitated through use of ImageJ’s Fibriltool.^81^ A minimum of 30 cells were analyzed per condition for each experiment. The calcium labeling images were analyzed using ImageJ where the Fura Red image was used to establish an appropriate threshold to minimize background signal while maximizing cell outlines. This threshold was then applied to the Fluo-4 image and the mean intensity value of each cell was recorded. The Fluo-4 intensity values were then divided by the corresponding Fura Red intensity values to provide the net calcium signal for each cell. The potassium labeling images were also analyzed using ImageJ where each image was thresholded to minimize background signal and maximize individual cell outlines. Once applied, the mean intensity values were recorded for each cell. MRTFA protein localization was quantitated manually and in a blinded manner. Mitochondrial aspect ratio, length and area were determined by mitochondria-analyzer FIJI macro.

## RESOURCE AVAILABILITY

### Lead contact

Information and requests for reagents may be directed to and will be fulfilled by the corresponding author, Dr. Ekrem Emrah Er, upon reasonable request.

### Materials availability

Requests for reagents will be fulfilled by the corresponding author upon reasonable request

### Data and code availability

All relevant data can be found within the article and its supplementary information and upon reasonable request from the corresponding author. RNA Sequencing data has been deposited under GSE273414 and GSE285078.

## ACKNOWLEDGEMENTS

We thank Dr. Morgan Huse and Dr. Maria Tello-Lafoz for the technical and scientific input during the preparation of this manuscript and Priya M Shah for her help with sample processing and for providing critical discussion. We thank Jiana Calixto and Dr. Paris Thomas of Equal Hope for their input as Er Lab’s community partner. We thank Dr. Andrius Kazlauskas and Marlen Gonzalez for their help with MitoTracker image analysis. We thank UIC Research Histology for their assistance with histological techniques and the Cardiovascular Research Core (CVRC) members Dr. Jiwang Chen and Dr. Samuel Lee for conducting Echocardiographic imaging and data analyses as part of CVRC services, respectively. This work was supported by NCI (MERIT AWARD R37CA269370 to EEE and R33-CA258012 to JR), NIA (R01AG044404, JCL and F31AG090005 to MAS), Concern Foundation (Conquer Cancer Now Award, EEE), American Cancer Society (RSG-24-1320151-01-MM) University of Illinois Cancer Center (Hope Leaders Award, EEE and AMG) and University of Illinois at Chicago (Physiology and Biophysics start-up funds to EEE).

## AUTHOR CONTRIBUTIONS

AMG and EEE conceived the project, designed the experiments, interpreted data and wrote the manuscript. AMG, MH, VP, KM, KJP, RRL, CCC, KL, SMW conducted experiments and interpreted the data. FH, ENM, JL provided critical feedback, technical and intellectual input into AFM experimental design and execution. MAS assisted with mRNA sequencing data parsing; mapping reads and differential gene expression. KMB performed electrophysiology experiments and interpreted data. EEE performed GSEA. IL, BMW and JR provided critical feedback and edited the manuscript.

---

**Figure S1.**
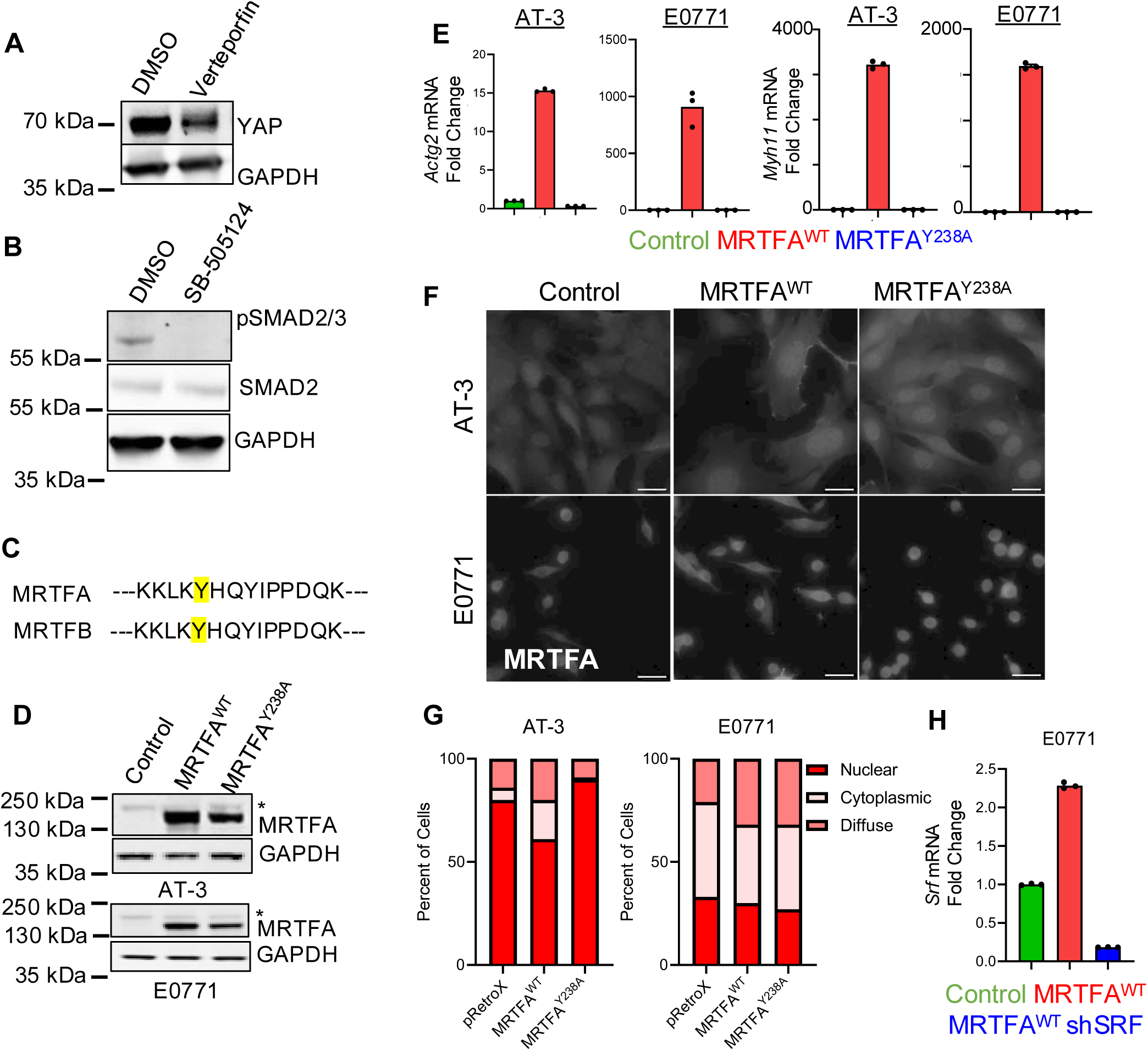
Related to Figure 1. (A) Immunoblot of YAP in E0771 MRTFA^WT^ cells treated with 10 µM verteporfin for 24 hours compared to DMSO control. GAPDH is used as loading control. (B) Immunoblot of SMAD phosphorylation in E0771 MRTFA^WT^ cells treated with 2.5ng/mL TGFß2 for 1 hour in the presence or absence of 2.5 µM SB-505124 pre-treatment. GAPDH is used as loading control. (C) MRTFA and MRTFB amino acid alignment around the conserved Y238 and Y305 residues, respectively. (D) Immunoblot of MRTFA overexpression in MRTFA^WT^ and MRTFA^Y238A^ cells compared to control. GAPDH is used as loading control. * indicates endogenous MRTFA. (E) Relative transcript levels of actin-related established MRTFA target genes *Actg2* and *Myh11* in MRTFA^WT^ and MRTFA^Y238A^ mutant overexpressing cell lines. Data are shown as fold change calculated from 3 technical replicates per sample and normalized to GAPDH. Each graph is representative of 3 independent experiments. (F) Representative immunofluorescent images depicting MRTFA protein localization in MRTFA^WT^ and MRTFA^Y238A^ mutant overexpressing cell lines. Scale bars = 50 µm. (G) Quantitation of MRTFA protein localization in MRTFA^WT^ and mutant overexpressing cell lines. AT-3: Control (pRetroX) = 298 cells, MRTFA^WT^ = 140 cells, MRTFA^Y238A^ = 169 cells; E0771: Control (pRetroX) = 185 cells, MRTFA^WT^ = 315 cells, MRTFA^Y238A^ = 232 cells. (H) Relative transcript levels of SRF in E0771 MRTFA^WT^ cell lines with/without SRF knockdown (shSRF). Data are shown as fold change calculated from 3 technical replicates per sample and normalized to GAPDH. Graph is representative of 3 independent experiments.

**Figure S2.**
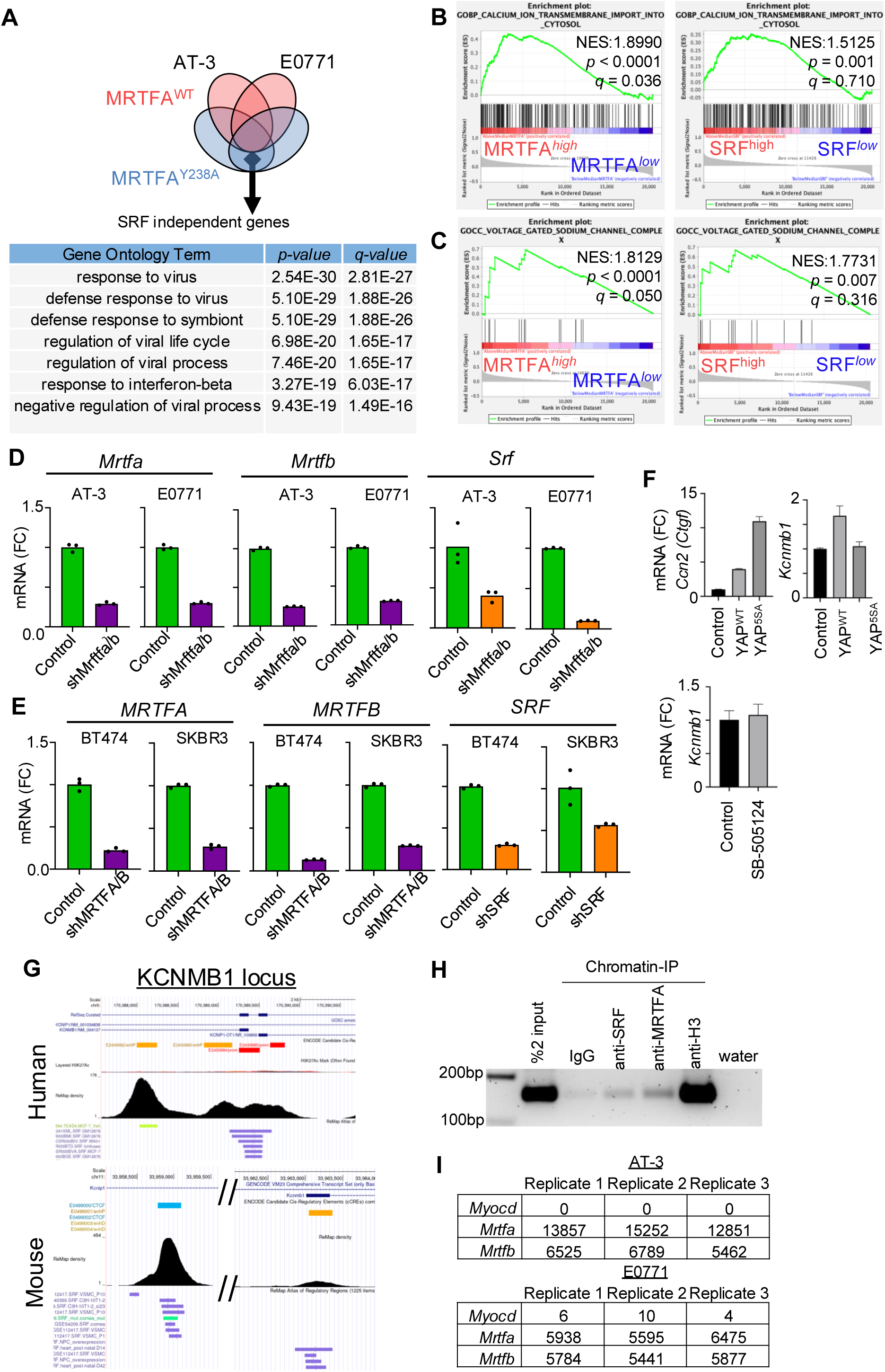
Related to Figure 2. (A) Venn diagram and gene ontology analyses of the SRF independent genes that are exclusively upregulated by MRTFA^Y238A^ in both AT-3 and E0771 cells to at least 2-fold or more. See Table S2 for full results. (B) Examples of statistically significant GSEA for MRTFA^high/low^ breast cancer patient tumors from TCGA. (C) Examples of statistically significant GSEA for SRF^high/low^ breast cancer patient tumors from TCGA. (D) Relative mRNA expression levels of MRTFA and MRTFB in shMRTFA/B and SRF in shSRF mouse cell lines. Data are shown as fold change calculated from 3 technical replicates per sample and normalized to GAPDH levels. Each graph is representative of 3 independent experiments. (E) Relative mRNA expression levels of MRTFA and MRTFB in shMRTFA/B and SRF in shSRF human cell lines. Data are shown as fold change calculated from 3 technical replicates per sample and normalized to GAPDH levels. Each graph is representative of 3 independent experiments. (F) Relative mRNA expression levels of canonical YAP target gene CCN2, which encodes connective tissue growth factor (CTGF), and KCNMB1 in AT-3 cells overexpressing wildtype YAP or hyperactive YAP^5SA^ mutant or treated for 24 hours with TGFβ inhibitor SB-505124. (G) KCNMB1 human and mouse genetic loci analyzed for SRF (purple tracks) binding by using University of California Santa Cruz Genome Browser data publicly available Chromatin immunoprecipitation sequencing data. Each purple solid line under the ReMap density map indicates an independent dataset where SRF was found to be associated with the locus. In the Encode cCREs track, red solid boxes indicate promoter regions, orange and yellow solid boxes indicate proximal and distal enhancers, respectively, and blue tracks indicate CTCF only binding regions. (H) Chromatin immunoprecipitation-PCR from AT-3 cells using KCNMB1 primers and the indicated antibodies against SRF, MRTFA, histone H3 (as a positive control) and IgG (negative control). Water was used as input material as a negative control for the PCR reaction. Image is representative of 3 independent experiments. (I) Raw counts from bulk RNA-seq data for *Myocd*, *Mrtfa*, and *Mrtfb* genes in AT-3 and E0771 cells.

**Figure S3.**
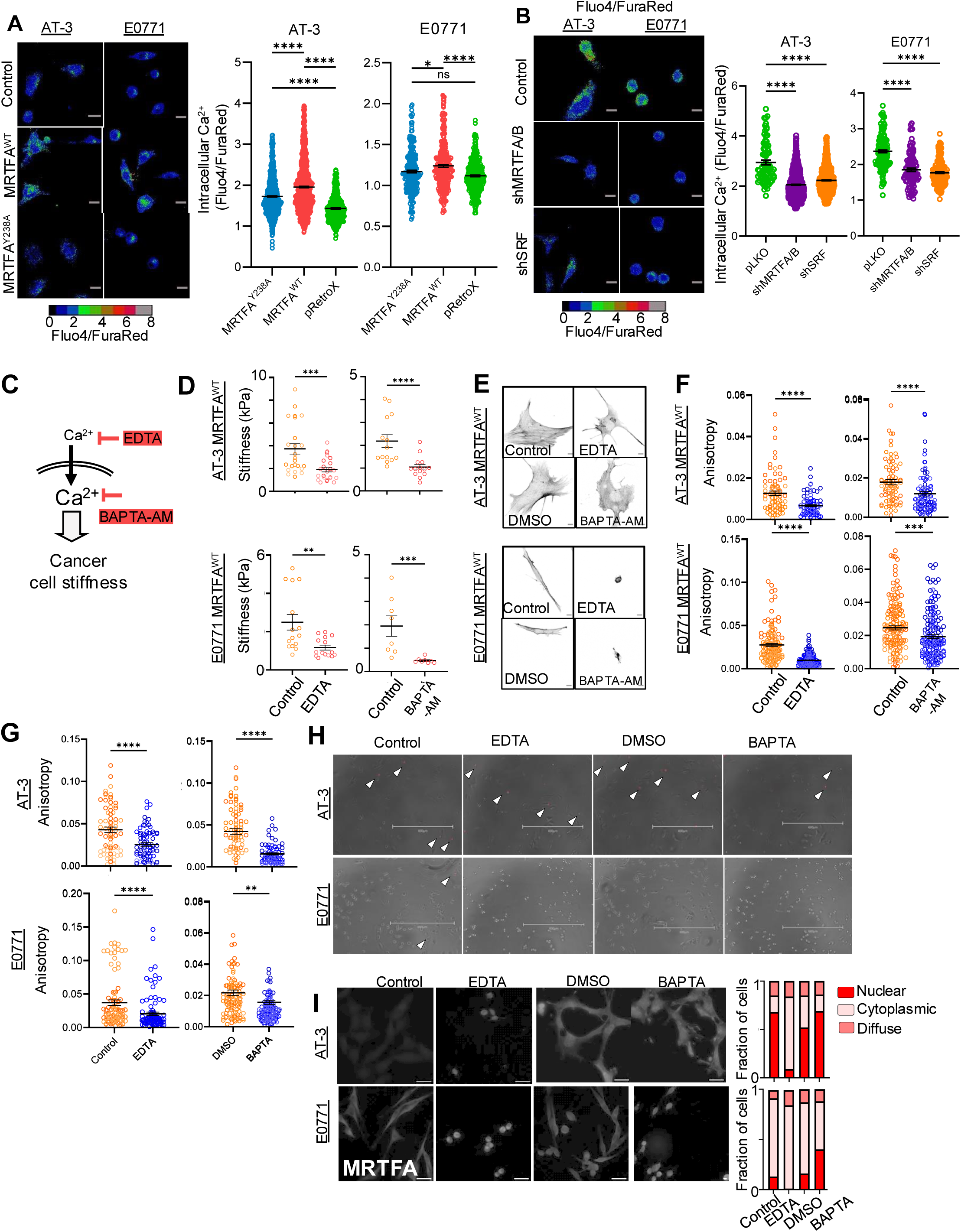
Related to Figure 3. (A-B) Left: Ratiometric images of representative cells loaded with Fluo-4 and FuraRed Ca^2+^ binding dyes. FuraRed signal indicates Ca^2+^-free dye loading and Fluo-4 indicates Ca^2+^-bound dye. Scale bar = 10 µm. Right: Graphs showing mean fluorescence intensity ratio of Fluo-4 and Fura Red. Each graph is representative of 3 independent experiments; AT-3: Control (pRetroX) = 415 cells, MRTFA^WT^ = 862 cells, MRTFA^Y238A^ = 670 cells, Control (pLKO) = 77 cells, shMRTFA/B = 924 cells, shSRF = 776 cells; E0771: Control (pRetroX) = 367 cells, MRTFA^WT^ = 303 cells, MRTFA^Y238A^ = 238 cells, Control (pLKO) = 140 cells, shMRTFA/B = 108 cells, shSRF = 190 cells; Kruskal-Wallace test with multiple comparisons. All error bars are mean±s.e.m. ns, not significant; *P<0.05; **P<0.01; ***P<0.001. (C) Schematic of different Ca^2+^ chelating agents their site of extra- and intra-cellular action. (D) AFM stiffness measurements of AT-3 and E0771 MRTFA^WT^ cells treated with EDTA or BAPTA-AM compared to DMSO control groups. Results are from 3 independent experiments combined. AT-3: Control (water as a control for EDTA) = 24 cells, EDTA = 23 cells, DMSO (Control for BAPTA-AM) = 22 cells, BAPTA-AM = 20 cells; E0771: Control (water) = 15 cells, EDTA=15 cells, DMSO (Control for BAPTA-AM) = 8 cells, BAPTA-AM = 9 cells; Mann-Whitney test. All error bars are mean±s.e.m. ns, not significant; *P<0.05; **P<0.01; ***P<0.001. (E) Confocal images of representative AT-3 and E0771 MRTFA^WT^ overexpressing cells treated with EDTA or BAPTA-AM compared to controls and stained with phalloidin (F-actin). Scale bar=10 µm. (F) Anisotropy measurements of cells in (E). Each graph is representative of 2 independent experiments per cell type and treatment. AT-3: Control (water as a control for EDTA) = 31 cells, EDTA=31 cells, DMSO (Control for BAPTA-AM) = 32 cells, BAPTA-AM = 35 cells; E0771: Control (water) = 46 cells, EDTA = 73 cells, DMSO (Control for BAPTA-AM) = 40 cells, BAPTA-AM = 43 cells; Mann-Whitney test. All error bars are mean±s.e.m. ns, not significant; *P<0.05; **P<0.01; ***P<0.001. (G) Anisotropy measurements of unmodified AT-3 and E0771 cells treated with EDTA or BAPTA, compared to respective controls. Each graph is representative of 2 independent experiments per cell type and treatment. AT-3: Control (water as a control for EDTA) = 77 cells, EDTA=74 cells, DMSO (Control for BAPTA-AM) = 76 cells, BAPTA-AM=89 cells; E0771: Control (water) = 93 cells, EDTA = 104 cells, DMSO (Control for BAPTA-AM) = 84 cells, BAPTA-AM = 95 cells; Mann-Whitney test. All error bars are mean±s.e.m. ns, not significant; *P<0.05; **P<0.01; ***P<0.001. (H) Representative images of AT-3 and E0771 cells treated with EDTA or BAPTA-AM and stained with cell death marker propidium iodide (PI, red) during live culture. Arrow heads point to PI^+^ cells that are never in high abundance under any condition. Images are representative of 3 independent experiments. Scale bar = 600 µm. (I) Left: Representative immunofluorescent images of MRTFA protein localization in cells treated with EDTA or BAPTA-AM, compared to respective controls. Scale bars= 50 µm. Right: Quantitation of MRTFA protein localization in cells as seen at left. Graphs are a compilation of 3 independent experiments. AT-3: Control (water as a control for EDTA) = 316 cells, EDTA = 69 cells, DMSO (Control for BAPTA-AM) = 357 cells, BAPTA-AM = 187 cells; E0771: Control (water) = 178 cells, EDTA = 72 cells, DMSO (Control for BAPTA-AM) = 234 cells, BAPTA-AM = 128 cells.

**Figure S4.**
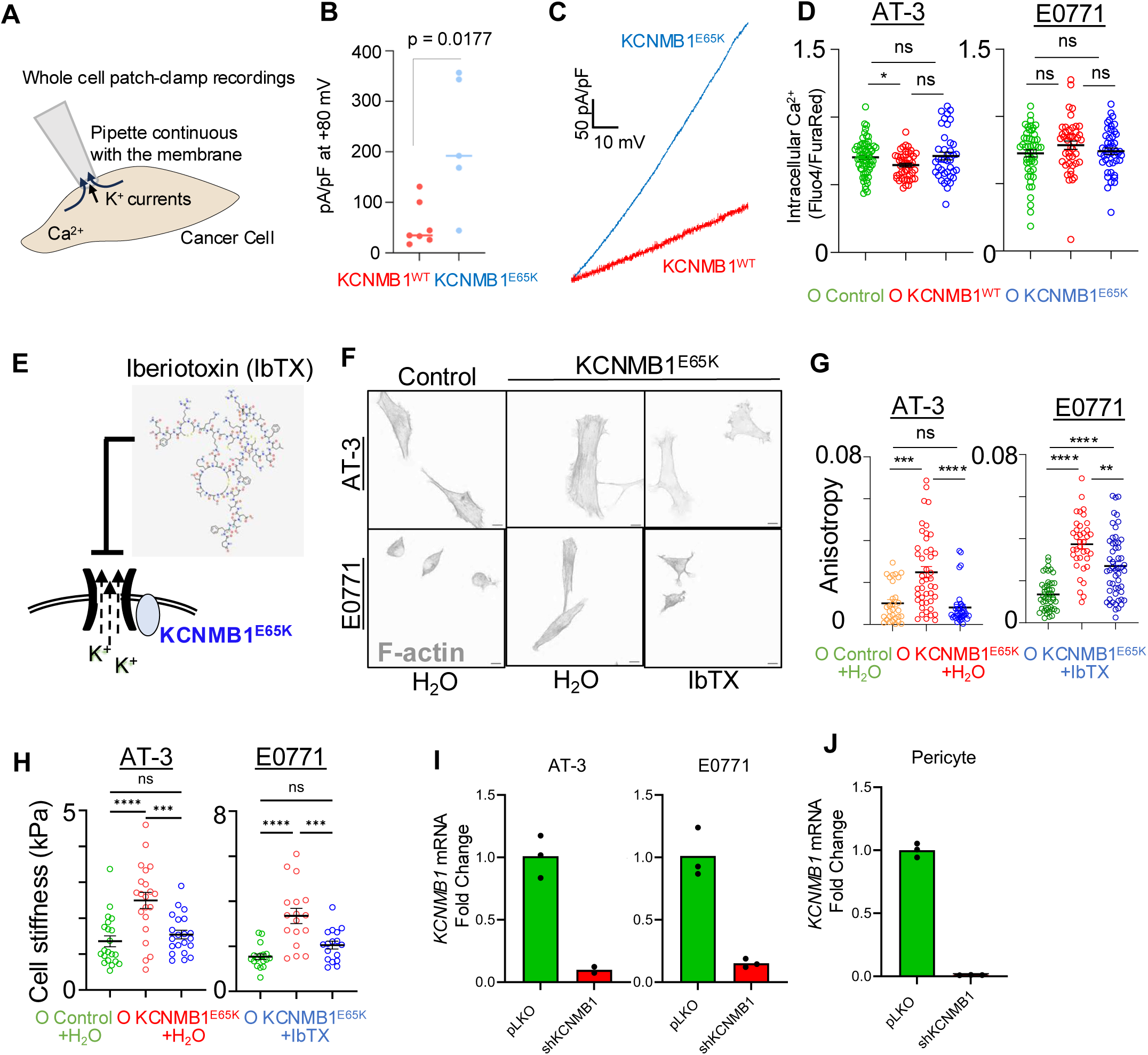
Related to Figure 4. (A) Whole patch clamp technique showing K^+^ current readings in cells that are stimulated with an intracellular Ca^2+^ as described in the methods. (B) Current density measurements at + 80 mV from AT3 cells overexpressing KCNMB1^WT^ (n = 7 cells) or KCNMB1^E65K^ mutant (n = 5 cells). *p* value calculated by non-parametric Mann-Whitney test. pA/pF: picoampere over picofarad. (C) Representative electrical current reading from an AT3 cell overexpressing KCNMB1^WT^ and an AT3 cell overexpressing KCNMB1^E65K^ mutant. Current readings were generated by OriginLab software. (D) Ratiometric intracellular calcium measurements performed in the indicated cell lines. Graphs are representative of 4 and 5 independent experiments using AT-3 and E0771 cell lines, respectively. n = 63 and 51 control cells, 46 and 44 KCNMB1^WT^ overexpressing cells and 40 and 46 KCNMB1^E65K^ overexpressing AT3 and E0771 cells, respectively. *p* values were calculated by Kruskal Wallis Test followed by Dunn’s test for multiple comparisons. *: *p* < 0.05, ns: *p* > 0.05. (E) Schematic of the experimental design that investigates the impact of IbTX, a non-cell permeable BK channel blocker in cells that overexpress the hyperactive KCNMB1^E65K^ variant. (F) Representative phalloidin staining images from AT-3 and E0771 cells expressing control vector or overexpressing KCNMB1^E65K^ mutant with or without BK channel inhibitor Iberiotoxin (IbTX) treatment. (G) Anisotropy measurements from cells depicted in (F). Graph is representative of 2 and 3 experiments for AT3 and E0771 cells. n = 30 and 44 control cells, 45 and 37 KCNMB1^E65K^ overexpressing cells and 37 and 53 KCNMB1^E65K^ + IbTX cells for AT3 and E0771 experiments, respectively. *p* values were calculated by Kruskal Wallis Test followed by Dunn’s test for multiple comparisons. ****: *p* < 0.0001, ***: *p* < 0.001, **: *p* < 0.01, *: *p* < 0.05, ns: *p* > 0.05. (H) Cell stiffness measurements in cells from F-G. Graph represents combined data from 3 independent experiments for each cell line. *p* values were calculated by Kruskal Wallis Test followed by Dunn’s test for multiple comparisons. ****: *p* < 0.0001, ***: *p* < 0.001, **: *p* < 0.01, *: *p* < 0.05, ns: *p* > 0.05. (I-J) Relative transcript levels of KCNMB1 in shKCNMB1 cell lines. Data are shown as fold change calculated from 3 technical replicates per sample and normalized to GAPDH. Each graph is representative of 3 independent experiments.

**Figure S5.**
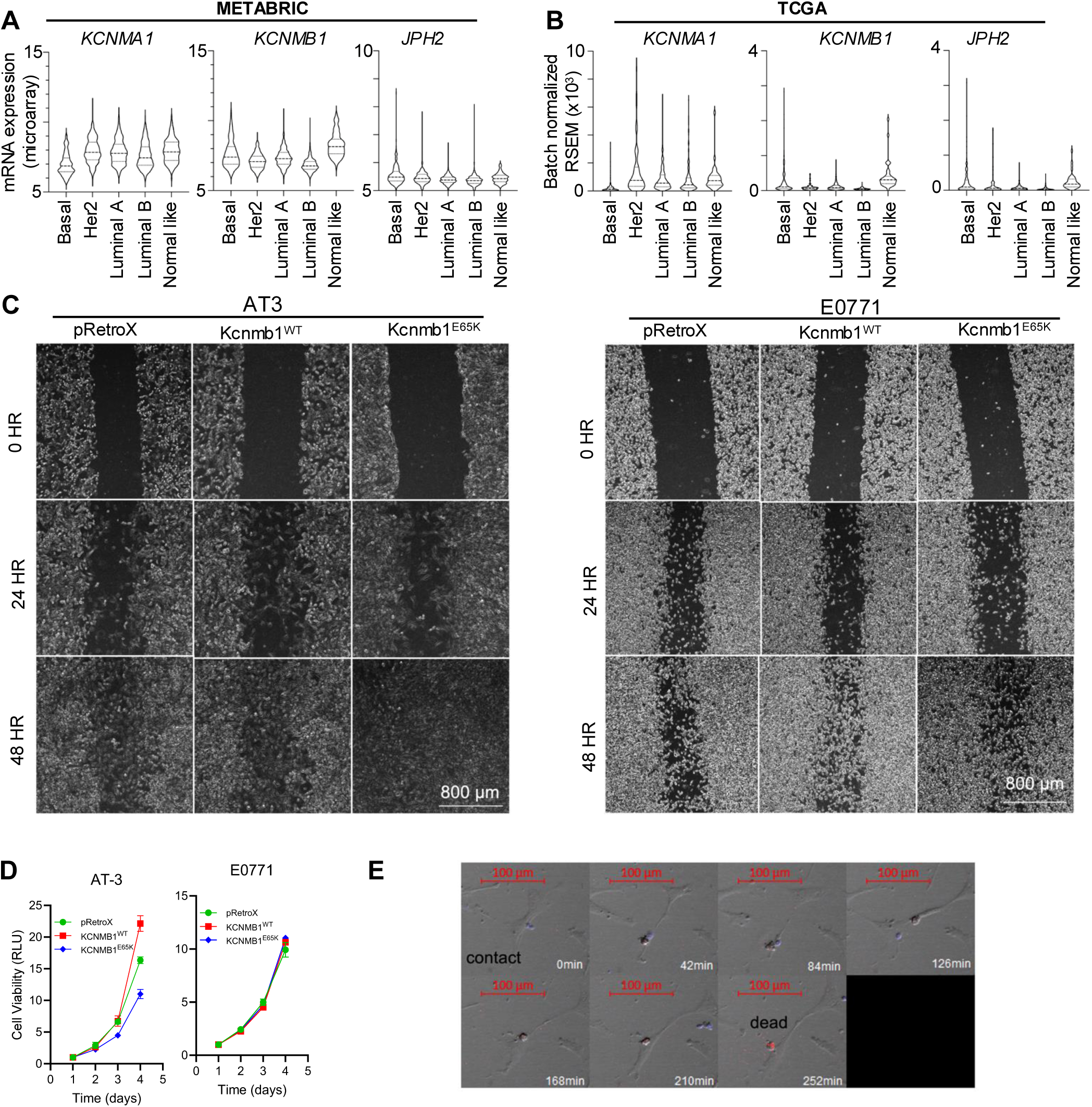
Related to Figure 5. (A – B) mRNA expression of the listed genes in the (A) METABRIC dataset and (B) TCGA PanCancer Atlas for Invasive Breast Cancer across the major PAM50 breast cancer subtypes in patients. See Table S5. (C) Wound healing assays in the indicated cell lines, imaged at the indicated time points. HR: hours. Images are representative of 3 independent experiments. Scale bars = 800 µm. (D) Relative cell viability in the indicated cell lines at the indicated at the indicated time points normalized to the Day 1 (24 hours) time point. Graph is compiled from 3 and 2 independent experiments performed in triplicate for AT3 and E0771 cells, respectively. Error bars: s.e.m. (E) Representative image for assessment of cell death as judged by membrane blebbing and eventual PI inclusion after cancer cell (bright field) is contacted by a natural killer cell (blue).

**Figure S6.**
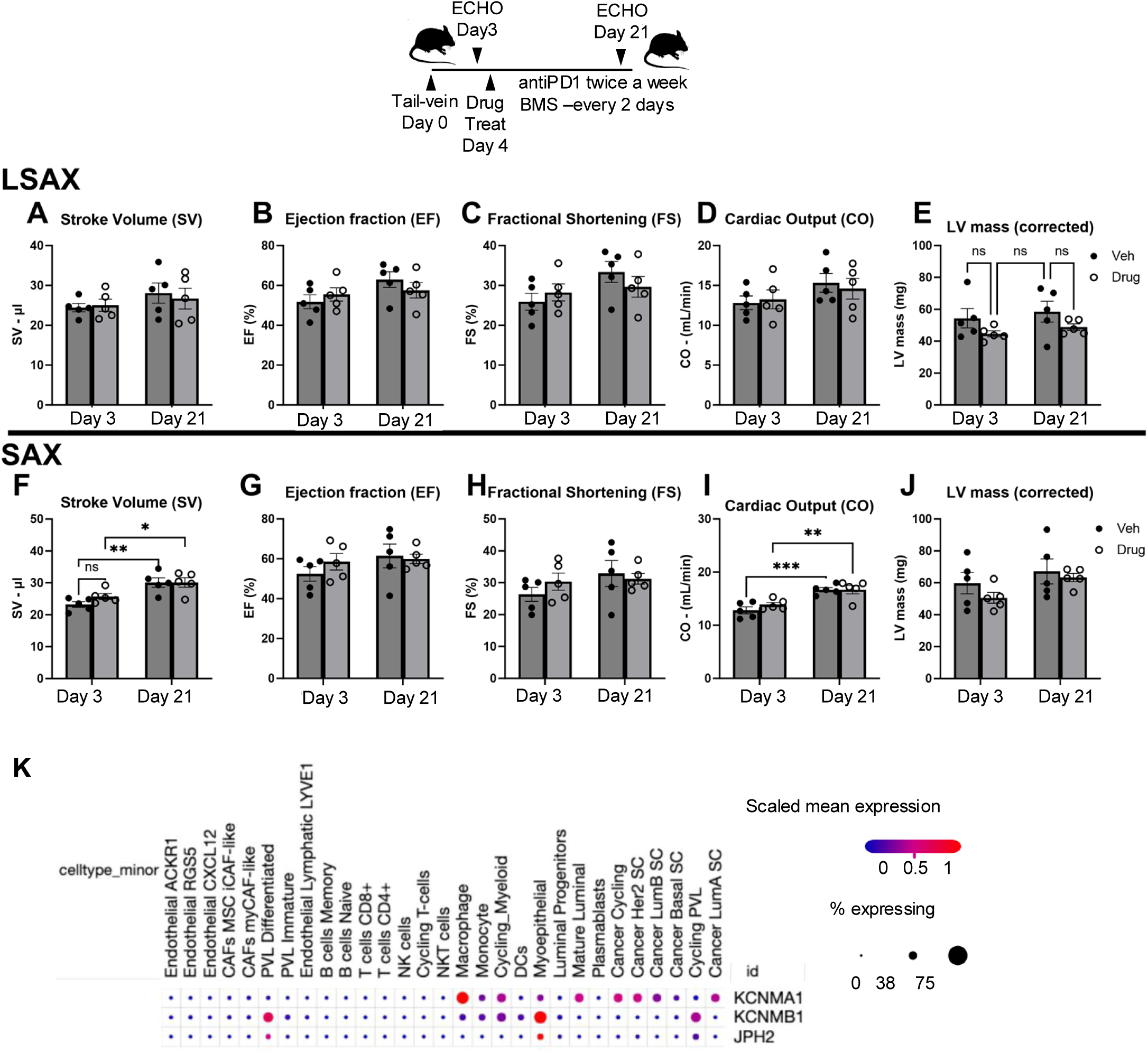
Related to Figure 6. (A-J) Assessment of Left ventricular systolic function and mass 3 days and 21 days after tail-vein injection of mammary tumor cells in vehicle/IgG control treated and antiPD1 and BMS-204352 treated female mice. n = 5 mice per group M-mode images of parasternal long axis (LSAX) and short axis (SAX) were analyzed for stroke volume (A, F), ejection fraction (B, G), fractional shortening (C, H), cardiac output (D, I) and left ventricular mass (E, J). Data presented as mean ± standard error of the mean with two-way ANOVA with posthoc TukeyHSD analyses; ** - P value <0.001 *** - P value = 0.0001. (K) KCNMA1, KCNMB1 and JPH2 expression in the cells of the breast tumor microenvironment accessed from Broad Institute’s Single Cell Portal.

**Figure S7.**
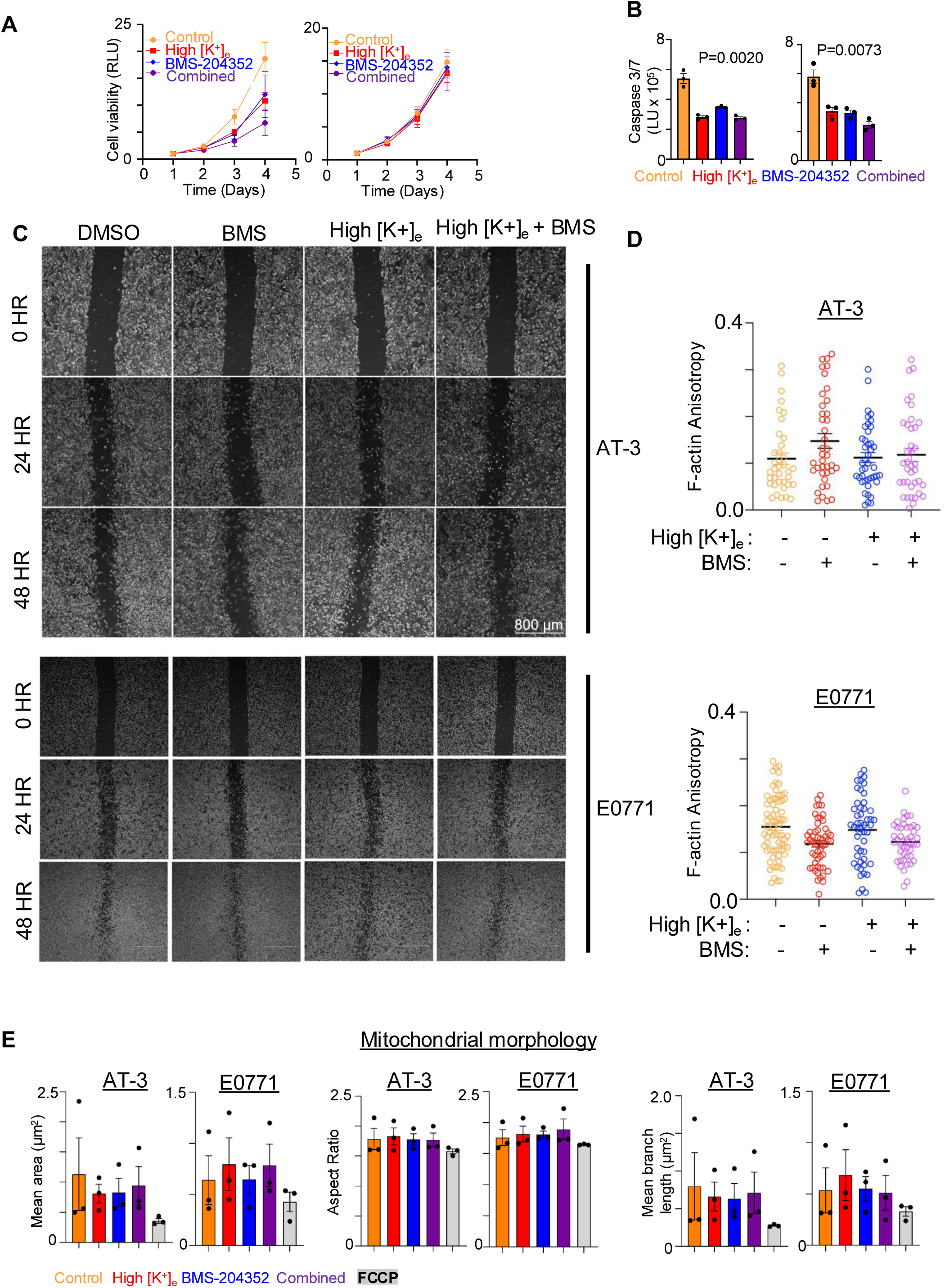
Related to Figure 7. (A) Cancer cell viability curves over the course of 4 days post-treatment with high [K^+^]_e_ (45 mM KCl) and 10 µM BMS-204352. Relative luminescence units (RLU) are normalized to untreated Day 1 samples. All error bars are mean±s.e.m. (B) Caspase 3/7 activity in cancer cells treated with with high [K^+^]_e_ (45 mM KCl) and 10 µM BMS- 204352 after 24 hrs. Graphs are representative of 3 independent experiments. All error bars are mean ± s.e.m. (C) Wound healing assays in the indicated cell lines, imaged at the indicated time points. HR: hours. Images are representative of 3 independent experiments. Scale bars = 800 µm. (D) F-actin anisotropy measurements of cells treated with the indicated agents. Data are representative of 3 independent experiments. (E) Mitochondrial morphology assessment in cells that were treated with the indicated agents. Carbonyl cyanide p-trifluoromethoxyphenylhydrazone (FCCP) was used as a positive control to disrupt mitochondrial morphology. Graph combines mean cellular values from three independent experiments.

